# SenCat: Cataloging human cell senescence through multiomic profiling of multiple senescent primary cell types

**DOI:** 10.64898/2026.02.05.703986

**Authors:** Carlos Anerillas, Gisela Altés, Katarína Grešová, Dimitrios Tsitsipatis, Krystyna Mazan-Mamczarz, Reema Banarjee, Ana S.G. Cunningham, Martin Salamini-Montemurri, Jen-Hao Yang, Rachel Munk, Martina Rossi, Yulan Piao, Bradley Olinger, Quinn Strassheim, Jennifer L. Martindale, Jinshui Fan, Chang-Yi Cui, Supriyo De, Delaney V. Rutherford, Ying Hao, Ziyi Li, Jessica Roberts, Yue Andy Qi, Kotb Abdelmohsen, Rafael de Cabo, Allison B. Herman, Manolis Maragkakis, Nathan Basisty, Myriam Gorospe

## Abstract

There is an urgent need to comprehensively catalog senescence markers across cell types in an organism in order to characterize ‘senotypes’ and senescent cell heterogeneity. Here, we profiled the transcriptomes and proteomes in 14 different primary human cell types undergoing over 30 senescence paradigms to create a senescence catalog we termed ‘SenCat’. We found that, while senescent cells from all primary tissue types did not share a single unique marker, they did activate shared specific metabolic and damage-response pathways implicated in tissue repair. Machine learning analysis of the SenCat transcriptomic and proteomic datasets successfully identified independent sets of senescent human cells, and senescent-like cells in mouse lung and kidney. In sum, SenCat represents a much-needed resource to identify senescent cells across tissues in the body.

**HIGHLIGHTS:**

- Identifying senescent cells in organisms in vivo remains a challenge
- We created SenCat: a catalog transcriptomes and proteomes of senescent primary cells
- Machine learning (ML) analysis of SenCat identified robust senescence scores
- ML-derived senescence scores uncovered senescent-like cell dynamics in vivo

## INTRODUCTION

Cellular senescence is a state of persistent cell cycle arrest triggered by sublethal damage, often accompanied by the secretion of bioactive molecules collectively known as the senescence-associated secretory phenotype (SASP). Through SASP factors, senescent cells communicate with their surroundings, contributing to physiological processes such as tissue regeneration, wound healing, and embryonic development^1,2^ However, when senescent cells accumulate abnormally, they can disrupt tissue homeostasis and promote fibrosis, chronic inflammation, and age-related pathologies.^3^ Given these dual roles in health and disease, there is growing urgency to comprehensively characterize senescent cells.^1,4,5^

Identifying and mapping senescent cells in tissues and organs has been a major challenge in the field^6,7^ and a major goal of the SenNet consortium.^5,8^ A key obstacle in senescence research is the inherent heterogeneity of senescent cells. Senescent phenotypes vary depending on the cell type, senescence-inducing stimulus, microenvironment, and time since senescence induction, with distinct transcriptomic and proteomic profiles reported across models.^9–11^ This complexity has impeded efforts to define universal senescence markers or to systematically identify ‘senotypes’—functional subtypes of senescent cells implicated in specific physiological or pathological outcomes.^12,13^ The lack of reliable and generalizable markers has also hindered efforts to map senescent cells within tissues and organs, particularly in complex *in vivo* settings.

To address these challenges, we launched the Senescence Catalog (SenCat) project: a large-scale, multimodal effort to define the molecular landscape of senescence. We systematically profiled the transcriptomes and proteomes of over 30 senescence models across 14 primary human cell types representing diverse lineages and tissue origins. Using RNA sequencing-based transcriptomic analysis and mass spectrometry-based proteomic analysis, we sought to identify both shared and context-specific features of the senescent state induced by various stimuli, including replicative exhaustion, DNA damage, and oncogenic stress. Our analysis revealed that senescence-associated gene expression changes are highly dependent on both the cell type and the trigger of senescence. While no universal markers emerged, we made several key discoveries. First, combinations of newly identified markers outperformed canonical markers in identifying senescent cells across conditions. Second, although the specific molecules involved varied, many of the underlying pathways altered in senescence were conserved, including canonical regulators such as p53 and NF-κB, as well as less-recognized pathways like epithelial-to-mesenchymal transition (EMT). Third, we developed a machine learning (ML)-based framework that integrated transcriptomic and proteomic features to derive robust, weighted senescence scores that consistently distinguished senescent from non-senescent cells in culture, complementing and strengthening existing efforts to define the senescent phenotype, including SenMayo, hUSI, SenSig, and Senepy.^14–17^ We assessed the utility of SenCat in vivo by analyzing single-nucleus (sn)RNA-seq data over time from lungs and kidneys of mice treated with the senescence-inducing agent doxorubicin. SenCat-derived markers captured dynamic, cell type-specific patterns of senescent cell accumulation, reinforcing the idea that senescence unfolds with distinct kinetics and phenotypic manifestations in different cell populations. A companion study highlights how the proteomic component of SenCat can be leveraged to develop clinically relevant, senotype-specific biomarker signatures in human plasma (Olinger et al. submitted)^18^. Together, these findings demonstrate that the SenCat resource not only improves our molecular understanding of senescence in culture but also enables the robust detection and characterization of senescent cells in complex tissues. The comprehensive transcriptomic and proteomic reference data in SenCat has created a robust framework to investigate senescence dynamics and functions in both experimental and physiologic contexts.

## RESULTS

### The SenCat project: profiling transcriptomes and proteomes in over 30 senescent cell models

To identify reliable and relevant markers of cell senescence, we embarked on the SenCat project, an effort aiming to elucidate and compare the transcriptomes and proteomes of senescent cells from a wide range of primary human cell types and lineages (**Figure 1A**). We included two fibroblast types (WI-38 from lung, BJ from skin), two epithelial cell types (HSAEC from lung, HEKn from skin), two endothelial cell types (HCAEC from artery, HUVEC from vein), as well as bone marrow mesenchymal stem cells (BMMSC), vascular smooth muscle cells (HVSMC), skeletal myoblasts (HSKM), peripheral blood mononuclear cells (PBMC), preadipocytes (PreAdipo), osteoblasts (NHO), astrocytes (NHA), and melanocytes (HEMn). We optimized and triggered cellular senescence in all of these cell types by different means (CTIS, chemotherapy-induced senescence; IRIS, ionizing radiation-induced senescence; OSIS, oxidative stress-induced senescence; and OIS, oncogene-induced senescence) following the doses and times indicated (STAR Methods); all comparisons were made to proliferating (P) or empty vector (EV) controls. The senescent state was confirmed by evaluating multiple senescence markers or phenotypes, consistent with guidelines from the SenNet consortium.^8^ First, growth arrest of the different cell types following senescence-triggering treatments, with a marked reduction in bromodeoxyuridine (BrdU) incorporation (**Figure S1**); we set the strict requirement that minimal or no cell death was seen to avoid noise from other forms of damage response such as apoptosis.^19–21^ Second, the treatments induced senescence-associated beta-galactosidase (SA-β-gal) activity in all senescence models, even if with different intensities and dynamics (**Figure S2**). Third, the treatments altered the levels of traditional senescence markers, *p16*, *p21*, *IL6*, *GDF15*, and *LMNB1* mRNAs. While multiple aspects of the senescence phenotype were induced in each model, a select subset of traditional senescence markers were not robustly associated with the implementation of this phenotype in all models (**Figures 1B** and **S3**), underscoring the great variability existing across different senescence models, and the need to find reliable and consistent senescence markers beyond those limited markers currently available.

**Figure 1.**
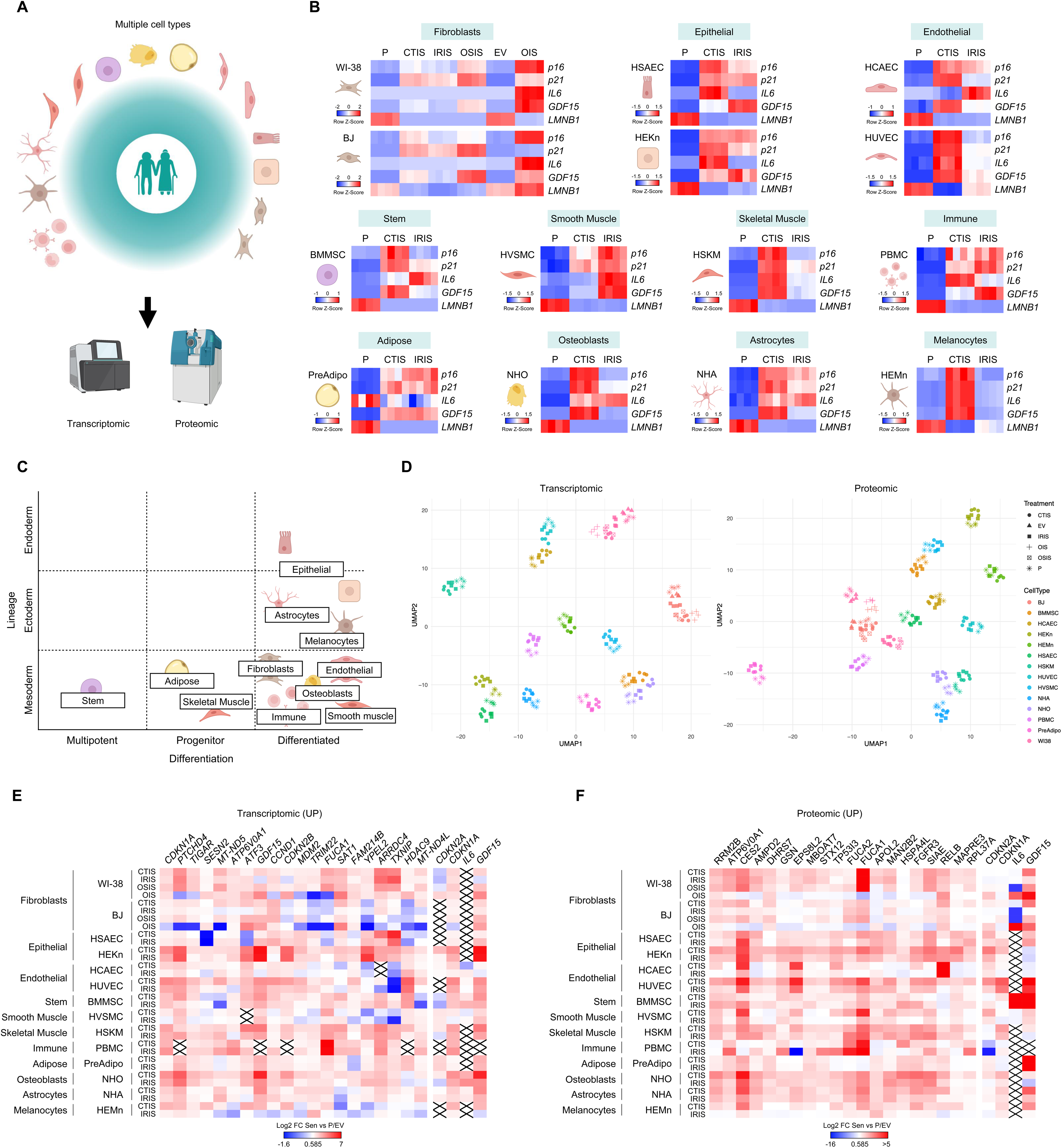
The SenCat project: transcriptomic and proteomic profiles of >30 senescence models. (A) Overview of the SenCat project: 14 cell types were subjected to senescence-inducing treatments followed by transcriptomic and proteomic profiling. (B) Heatmap representations of row Z-scores of the levels of canonical senescence-associated mRNAs measured by RT-qPCR analysis among the different cell types and senescence triggers. More details in **Figure S3**. (C) Distribution of the cell types in the SenCat project, depending on lineage and differentiation status. More details in **Figure S4A**. (D) UMAP clustering of the different cell types included in the SenCat project following transcriptomic (*left*) and proteomic (*right*) analyses. Proliferating controls are also included. Different colors indicate different cell types, and different shapes represent different senescence triggers. (E,F) Heatmaps displaying the Log2 FC in every senescence model (compared to their respective controls) for each of the top 20 most shared upregulated RNAs (E) and proteins (F) among senescent cell types observed. Canonical senescence markers *CDKN2A*, *CDKN1A*, *IL6*, and *GDF15* mRNAs (and encoded proteins) are displayed on the far right in both heatmaps. Crossed rectangles indicate no detection above the established threshold (Methods) of the indicated marker in a particular model. See also **Figures S1-S7**.

We compared the proteomic and transcriptomic profiles of the different senescent cells relative to their respective proliferating controls. **Figure 1C** shows all the cell types in the catalog by cell lineage and differentiation stage. As seen in **Figures 1D** and E, the most relevant category for classifying these cells by similarity in transcriptomic or proteomic expression profiles appears to be cell type, rather than tissue/lineage origin or differentiation stage (see also Figures **S4** and **S5**).

Importantly, we did not identify any universally shared senescence marker—RNA or protein (**Tables S1-6**). The lack of universal markers has been appreciated by different teams for some time^1,6^ and our comprehensive analysis confirms this notion. Analyzing the extent to which markers were shared within each cell type among the different senescence triggers revealed that, by increasing the numbers of transcriptomes or proteomes from senescence paradigms for a given cell type, the number of shared markers declined (**Figures S6**A-D) shortening the lists of shared markers. Even when the analysis focused on direct DNA-damaging agents (e.g., chemotherapy and IR), only ∼one-half of increased or reduced markers overlapped (RNA or protein; **Figures S6**A-D). Together, these findings indicate that senescence markers are not universal, and instead are dependent on trigger and cell type.

We also analyzed each cell type’s overlap between the transcriptomic and proteomic senescence markers. Despite including all senescence markers shared by at least 4 cell types, no cell type showed an overlap significantly higher than 20% (**Figures S6**E-G), indicating little concordance between transcriptomic and proteomic markers of senescence, and reinforcing the idea that cell senescence is complex and regulated at multiple levels affecting RNA and protein abundance.

After finding no single universal RNA or protein marker of cell senescence, we proceeded to analyze the most conserved sets of molecules across all the cell systems and triggers. As shown for increasing putative RNA and protein (**Figure 1**E and F) markers, as well as for declining putative RNA and protein (**Figure S7**A and B) markers, no marker changed consistently and significantly above or below 0.585 Log2 fold change (FC) (equivalent to 1.5 FC in natural scale) in all models of senescence (**Tables S3-S6**). Canonical markers often increasing in senescence, such as *CDKN1A* (*p21*) and *GDF15* mRNAs, were among the top 20 most shared markers upregulated by all senescence models in our analysis (**Figure 1E**), but, remarkably, CDKN1A/p21 and GDF15 proteins were not among the top 20 shared elevated markers (**Figure 1F**). Among the downregulated markers, however, the widely recognized markers *LMNB1* mRNA and LMNB1 protein were consistently reduced (**Figures 7A,B**).

We then compared the expression levels of the 20 identified top increased markers (RNA and protein) to those of canonical markers by evaluating average Log2 FC values for all the senescence models. As shown, the top 20 elevated RNA and protein markers were generally better at indicating senescence than the canonical markers (**Figure S7**C and D); of note, some of these new markers are also canonical (namely *CDKN1A* and *GDF15* mRNAs). Taken together, our analysis strongly supports the idea that no single marker is universally altered in all senescence, highlighting the need to explore alternative strategies to reliably identify senescent cells.

### Pathway analysis characterizes senescent cells as damage-responding cells that engage in tissue repair processes and metabolic reprogramming

In the absence of specific universal markers, we hypothesized that different gene expression programs may instead converge on shared cellular pathways differentially represented in senescent cells. We assessed pathways from the senescent cell transcriptomes in two ways: by including all the significantly changed (elevated and reduced) RNAs from each cell type separately and by including all the markers shared by at least 4 cell types (**Tables S3-6**). Analysis of the cell type-specific markers using the “Hallmark” pathway database^22^ pointed to several robustly enhanced pathways in almost every cell type: p53, EMT, NF-κB, apoptosis, and hypoxia (**Figure 2A**). Analysis of RNA markers shared by at least 4 cell types revealed the same pathways among the top 5 (**Figure 2B**, top-and bottom-left graphs), further suggesting that the core pathways associated with cell senescence were conserved regardless of cell type. Analysis directed at “GO Cellular Components” suggested that most pathways activated in senescence were related to lysosome function (**Figure 2B**, top-and bottom-right graphs). Meta-analysis of all the senescence-associated pathways grouped by the general processes to which they belong (**Figure 2C**) revealed that three pathways were particularly represented: metabolic reprogramming (blue oval), tissue remodeling (green), and response to damage (orange). Analysis of the reduced transcriptomic markers indicated that cell cycle progression-related pathways were inhibited in senescent cells (**Figure S8**A, **S6**B top-and bottom-left graphs), nuclear functions were diminished (**Figure S8**B, top-and bottom-right graphs), and there was broad inhibition of DNA replication and metabolism, and RNA metabolism and processing (**Figure S8**C).

**Figure 2.**
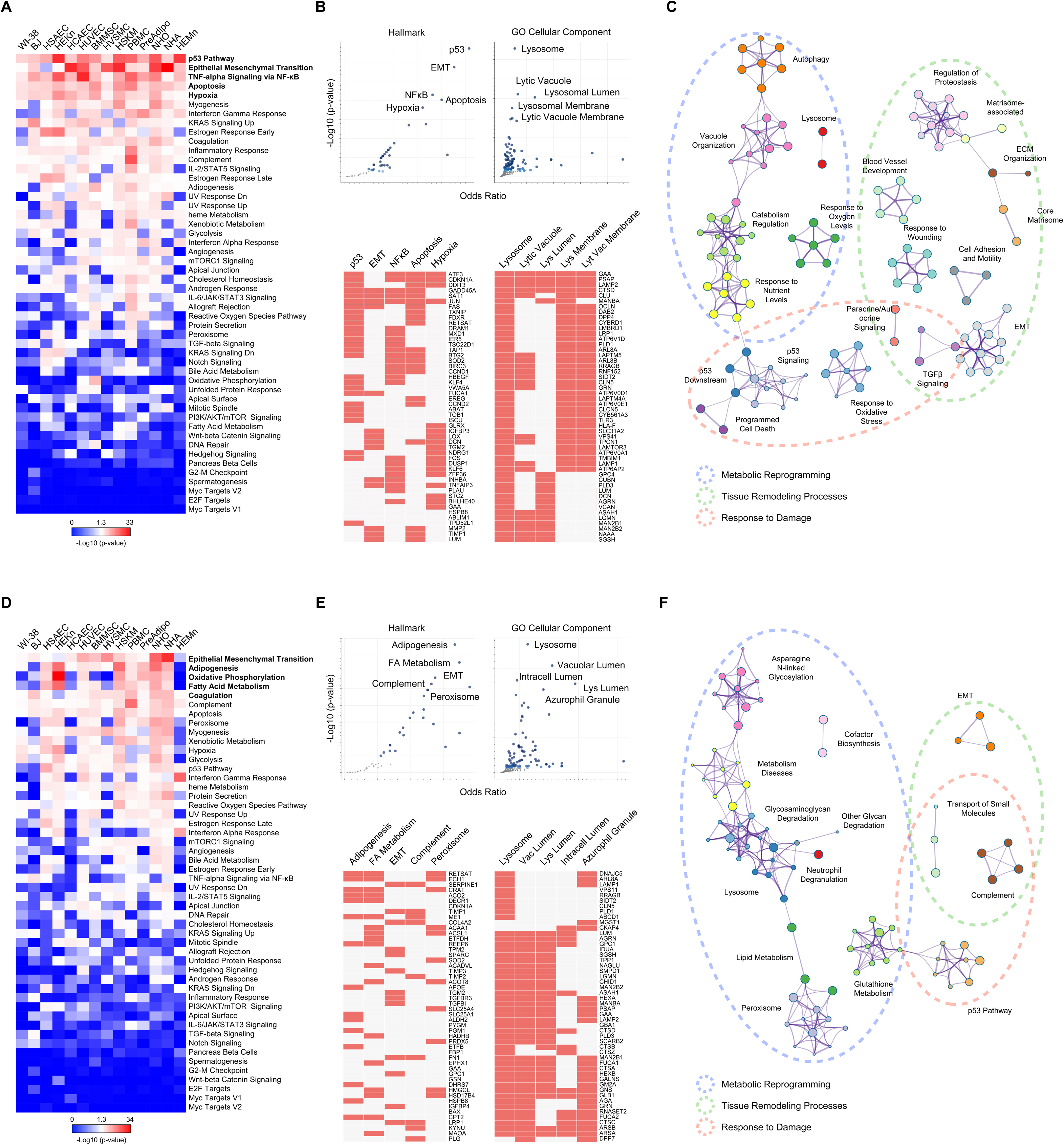
Senescent cells identified as damage-responding cells that engage in tissue repair processes while reprogramming specific metabolic routes (A) Heatmap displaying the association score (-Log10 p-value) of each cell type’s transcriptomic senescence markers with the indicated “Hallmark” pathways. (B) Dot plots (top) and marker-pathway association grids (bottom) performed with EnrichR show the association between the transcriptomic senescence markers shared by at least 4 cell types with the indicated “Hallmark” and “GO Cellular Component” pathways (top dot plots). Bottom grids display 50 markers associated with a specific pathway from those categories. (C) Metascape plot (including pathway categories only in the analysis) showing each of the pathways associated with the transcriptomic senescence markers shared by at least 4 cell types, clustered by common pathways and broader processes. (D-F) Analyses in D, E, and F are the same as in (A-C), but carried out using the proteomic data instead. See also **Figure S8**.

Turning to the proteomes upregulated with senescence, the only pathway conserved among those emerging from the transcriptomic analysis was the ‘EMT pathway’ (**Figure 2D** and E, top-and bottom-left graphs), in keeping with recent reports on the relevance of this pathway for senescent cells.^19–21,23^ Metabolic pathways (“adipogenesis”, “oxidative phosphorylation”, “fatty acid metabolism”), and “coagulation”, were also associated with cell senescence (**Figure 2D**), although with higher variability among cell types when compared to the transcriptomic pathway analysis. When considering proteomic markers shared by at least 4 cell types, the results were similar except for the inclusion of “peroxisome” as an enriched “Hallmark” pathway (**Figure 2E**, top-and bottom-left graphs); for the “GO Cellular Components” analysis, most lysosome-related pathways matched the transcriptomic results (**Figure 2E**, top-and bottom-right graphs). The general processes described for the transcriptomic analysis (“metabolic reprogramming”, “tissue remodeling”, and “response to damage”) were also conserved in the proteomic analysis (**Figure 2F**), although the internal composition of each of these categories varied somewhat. Analysis of the proteomic markers declining with senescence matched the transcriptomic results, with a significant reduction in processes related to cell cycle, the nucleus, DNA, and RNA (**Figure S8**D-F).

Together, the increased transcriptomic markers indicate a tilt towards damage response-related pathways, and the increased proteomic markers towards tissue repair and metabolic reprogramming pathways. The reduced transcriptomic and proteomic markers consistently showed declines in processes pertaining to nucleus function, cell cycle, and DNA and RNA processing. In sum, although senescent cells do not show strongly conserved individual changes in transcriptomes and proteomes, they do display a robust conservation in the pathways in which such senescence-associated RNAs and proteins participate. A further conclusion is to consider tailoring senotherapeutic strategies towards these conserved pathways rather than targeting a single transcript or protein.

### Machine learning (ML)-based identification of reliable senescence marker scores

To identify robust and generalizable markers of cellular senescence, we adopted a machine learning (ML) approach based on the SenCat transcriptomic and proteomic datasets. The strategy (**Figure 3A**) involved training penalized logistic regression models on RNA or protein data to classify senescent versus non-senescent samples. Each model was trained using a leave-one-cell-type-out cross-validation strategy, ensuring that markers were not biased toward any single lineage. Features consistently identified as predictive across all models were assigned final weights, and a senescence score was computed as the weighted sum of normalized expression values for each sample. Notably, *MX1*, *CXCL8*, and *GDF15* mRNAs, and FDXR, VDAC2 and TAGLN proteins showed strong positive coefficients, while *TOP2A* mRNA and some mRNAs encoding histones, along with SERPINH1, STMN1 and several histones, showed strong negative coefficients. The resulting transcriptomic and proteomic scores (**Figure 3B**) accurately discriminated senescent cells from controls across a wide panel of cell types and senescence triggers (**Figure 3C** and D). Importantly, while traditional senescence markers such as p16, p21, IL6, GDF15, and LMNB1 showed variable performance depending on cell lineage and condition, the ML-derived senescence scores consistently and robustly stratified senescent versus proliferating cells across the SenCat samples profiled (**Figure 3C**).

**Figure 3.**
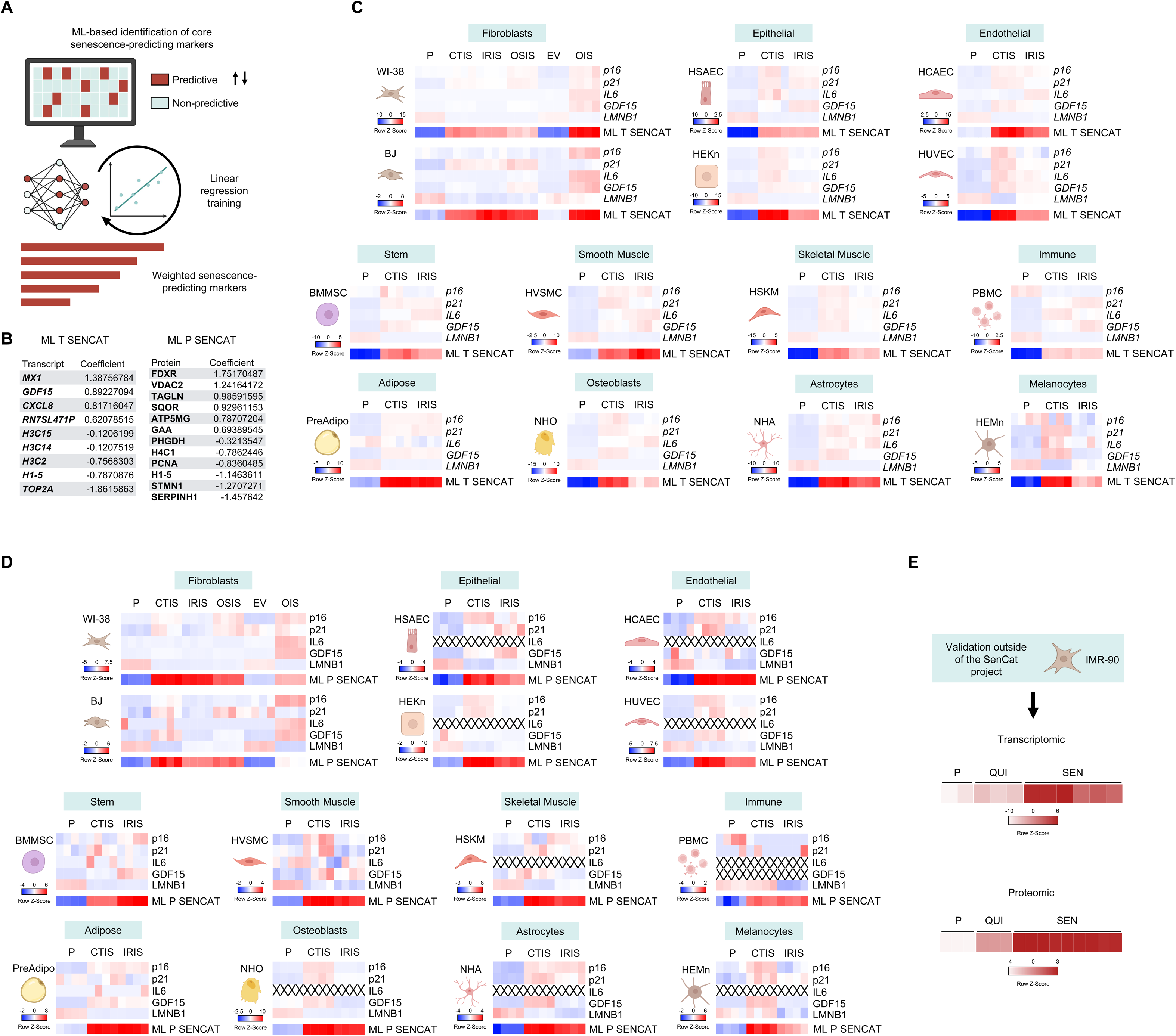
Machine learning (ML)-guided identification and validation of transcriptomic and proteomic senescence signatures across cell types and conditions. (A) Overview of the strategy used to identify core senescence markers using ML. Linear regression models were trained on transcriptomic and proteomic datasets from SenCat, and features consistently predictive across all models were assigned average coefficients (weights). (B) The resulting weighted transcriptomic (ML T SENCAT) and proteomic (ML P SENCAT) sets define a senescence score derived from a linear combination of marker expression levels and their respective ML-derived weights. (C) Heatmaps of traditional senescence markers (p16, p21, IL6, GDF15, LMNB1) and ML-derived transcriptomic senescence scores (ML T SENCAT) across 14 different human cell types subjected to senescence-inducing treatments (CTIS, IRIS, OSIS, OIS) or controls (P or EV) from the transcriptomic data. (D) Equivalent heatmaps displaying canonical senescence markers and ML-derived proteomic senescence scores (ML P SENCAT) as in (C), but obtained from the proteomic data. (E) Validation of ML-derived senescence scores on independent transcriptomic (top heatmap) and proteomic (bottom heatmap) datasets (IMR-90 lung fibroblasts external to the SenCat dataset). Quiescent (QUI), proliferating (P), and senescent (SEN) states were analyzed in these datasets. See also **Figures S8,9**.

To assess the generalizability of the ML-derived senescence signatures, we applied the transcriptomic and proteomic scoring systems to an independent dataset, IMR-90 lung fibroblasts treated to induce either quiescence or senescence (**Figure 3E**). Using transcriptomic and proteomic data, ML scores successfully distinguished senescent from non-senescent states, confirming that the signatures are not overfitted to the original training dataset and were applicable to new biological contexts. This finding was of vital importance, as it ensured SenCat datasets distinguished quiescent from senescent cells. Together, these results highlight the utility of an ML-guided strategy for identifying lineage-independent, multimodal markers of senescence.

### Validation of ML-derived senescence markers in vivo Across Senescence Models

To test if the ML-derived senescence signature (SenCat ML all) was capable of identifying senescent cells in physiologic settings in vivo, we generated single-nucleus (sn)RNA-seq data from lungs and kidneys of mice undergoing systemic senescence. As outlined in **Figure 4A**, we induced senescence through administration of doxorubicin (10 mg/kg), collecting lung and kidney samples at days 0, 1, 2, 4, 6, and 12 to capture the dynamics of senescence establishment; of note, we removed day 12 samples from the lung’s pool due to technical issues. Given the inherent limitations of snRNA-seq, including shallow coverage and a high-dropout rate, we applied the full ML-generated gene lists to maximize sensitivity (**Table S7**). Following clustering and cell type annotation (**Figures 4B** and C, **S11**A), we scored individual cells for senescence using AUCell.^24^ Benchmarking against existing senescence signatures (SenPy, SenSig, SenMayo, hUSI; **Table S8**), revealed superior performance by the ML-derived signature (SenCat ML all) in discriminating senescent from non-senescent cells across time points (**Figure S11**B). To rule out potential artifacts caused by cell-specific differences in sequencing depth, total UMI counts (nCount_RNA) were regressed from AUCell scores, obtaining similar results (**Figure S11**B). The need to use combinations of multiple senescence markers is even more obvious when testing the canonical markers of senescence, which are far less robust and reliable in their identification (**Figure S11**C, **S12**D).

**Figure 4.**
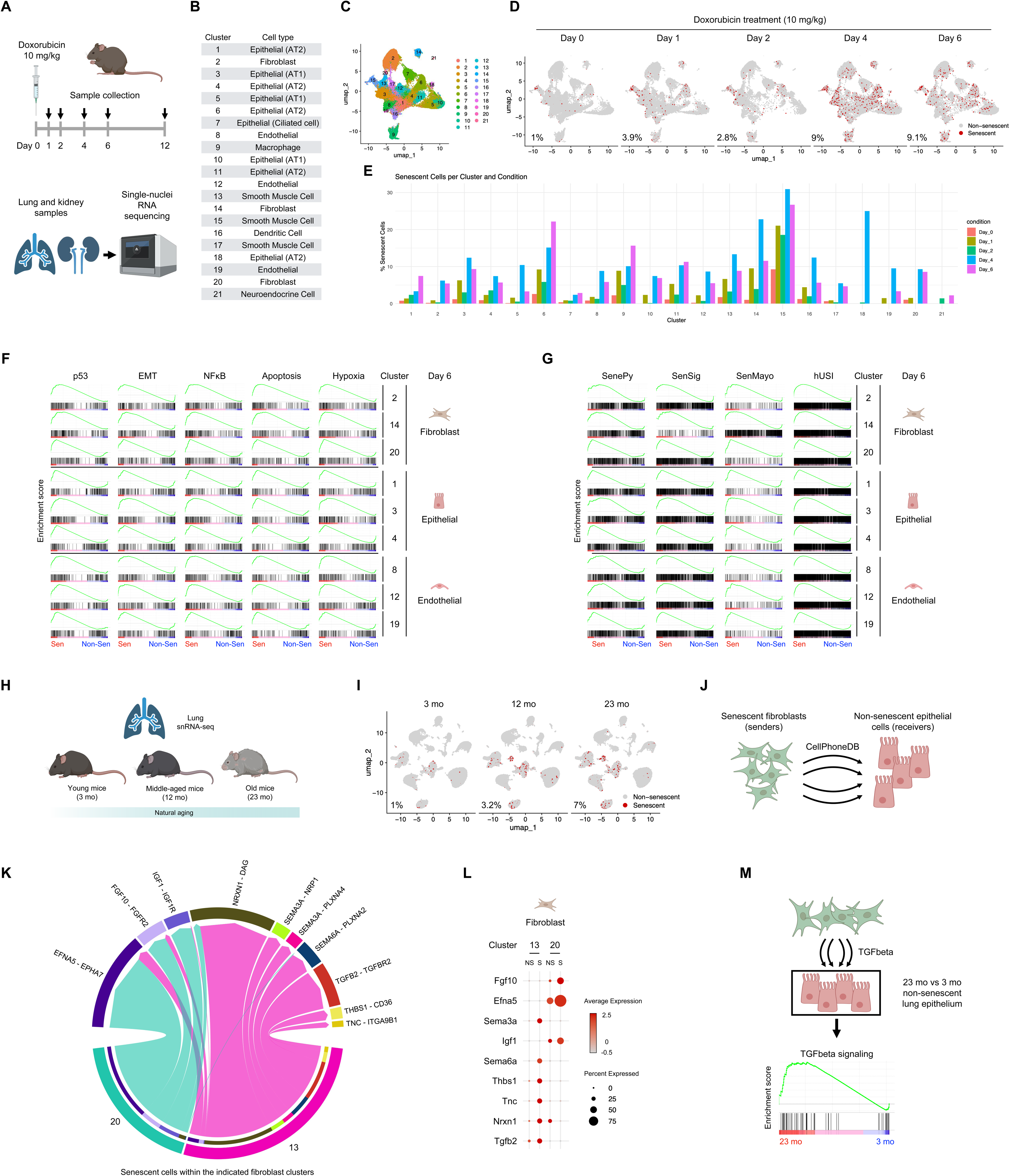
Identification of senescent cells in organs in vivo, leveraging ML-derived senescence marker signature. (A) Schematic of the experimental design for induction of systemic senescence with doxorubicin and sample collection across time points. (B) Cluster identities assigned to doxorubicin-treated lung samples profiled by snRNA-seq. (C) UMAP projection of lung samples colored by cluster identity. (D) UMAP projections of senescent cells in lung samples across time points. (E) Bar plot showing the number of senescent cells from lung per cluster and condition. (F,G) GSEA plots of gene set association scores for p53, EMT, NF-κB, Apoptosis, and Hypoxia hallmark pathways (F) and senescence signature lists SenePy, SeneSig, SenMayo, hUSI (G); in fibroblast, epithelial, and endothelial clusters on Day 6. (H) Schematic of the analysis pipeline applied to published aging lung snRNA-seq datasets. (I) UMAP projection of aging lung samples showing senescent cell distribution by age group. (J) Schematic of ligand-receptor inference analysis between senescent fibroblasts and non-senescent epithelial cells. (K) Chord diagram displaying ligand-receptor interactions between senescent fibroblasts (sender cells) and non-senescent epithelial cells (receiver cells) inferred through CellPhoneDB. (L) Dot plot showing expression of the specified ligands across fibroblast clusters at 23 months of age. (M) Schematic and GSEA plot explaining and displaying TGFβ signaling pathway increase during aging in non-senescent epithelial cells at 23 months versus 3 months. See also **Figures S11-13**.

We enhanced the performance of our approach by using the samples with minimal senescence (Day 0 samples) as a negative reference to refine the SenCat ML all list via an iterative ML approach (STAR Methods). Importantly, with this refined scoring approach we visualized the dynamics of senescent cell accumulation in lung at both global and cell type-specific levels (**Figure 4D** and E; **Figure S11**D). The same pipeline applied to kidney samples yielded similar dynamic changes in senescent cells (**Figure S12**A-H), underscoring the versatility of this strategy across organs.

We then investigated whether the transcriptomic programs associated with senescence observed in cultured cells persisted in live organs. We examined the GSEA association scores of the top 5 pathways associated with senescence (identified in **Figure 3**) across fibroblast, epithelial, and endothelial populations. Notably, within each of these lung cell lineages, senescent cells displayed increased representation of pathways including p53, EMT, NF-κB, apoptosis, and hypoxia, compared to non-senescent counterparts (**Figure 4F**). In addition, the ML-defined senescent populations were highly enriched for gene expression patterns captured by other senescence signatures,^14–17^ reinforcing the biological validity and generalizability of the ML-based classification (**Figure 4G**).

We then explored whether this same strategy could be used to dissect senescence dynamics during physiological aging. Using publicly available snRNA-seq data from lungs of young [3-month-old (mo)], middle-aged (12 mo), and old (23 mo) mice (**Figure 4H**), our refined senescence scoring pipeline uncovered an age-dependent increase in senescent cells (**Figure 4I**; **Figure S13**A-F), supporting the notion that senescent cells progressively accumulate in lung during aging.

Given the presumed role of senescent fibroblasts in driving epithelial dysfunction during aging, we next focused on fibroblast-to-epithelial cell communication. Using CellPhoneDB,^25^ we inferred ligand-receptor interactions between senescent fibroblasts and non-senescent epithelial cells in lungs from old mice (**Figure 4J**), an interaction long-suspected to be implicated in functional loss in the lung epithelium during aging. This analysis revealed numerous signaling axes potentially mediating this crosstalk (**Figure 4K**; **Figure S13**G); standing out among them was TGFB2 signaling, with pro-fibrotic and epithelium-remodeling roles (**Figure 4**L). In this regard, we observed a significant representation of the TGFβ pathway in non-senescent epithelial cells from 23-mo lungs compared to those from 3-mo (**Figure 4**M), consistent with enhanced paracrine signaling from increased numbers of senescent neighbors and with established features of epithelial aging.^21,23,26–29^

Together, these results indicate that the ML-derived senescence signature developed here can robustly identify senescent cells across tissues, time points, and senescence triggers in a live organ. Importantly, this approach performs well in both acutely induced and naturally occurring senescence, providing a high-resolution, generalizable framework for investigating senescence-associated processes across diverse biological contexts.

## DISCUSSION

We have carried out a comprehensive effort to characterize cellular senescence across multiple cell types and lineages. Our results underscore the complexity of senescence and the diversity of senescence-associated gene expression programs across cell types and triggers of senescence. A major challenge in the field has been to identify senescent cells in tissues and organs.^30–33^ Several studies have addressed this challenge using genetic and pharmacologic approaches to detect and clear senescent cells *in situ*,^34,35^ underscoring the interest in identifying senescent states within complex tissue environments. Recent technological advances, including single-cell RNA sequencing (scRNA-seq) and spatial transcriptomic analyses, offer unprecedented resolution for studying senescence in the context of the organism. Our results indicate that no single marker can universally identify senescent cells across all contexts, as previously proposed,^7^ and support the need to identify senotype-agnostic signatures. Instead, conserved biological programs (such as those involving p53, NF-κB, EMT, apoptosis, hypoxia, and lysosomal function) were increasingly represented across transcriptomic and proteomic datasets, regardless of the specific RNAs or proteins involved.

To address this limitation, we developed and implemented a machine learning-guided approach to identify lineage-independent transcriptomic and proteomic signatures of senescence. By training logistic regression models on the large and diverse senescence models in the SenCat dataset, we identified core RNAs and proteins that consistently distinguished senescent from non-senescent states across 14 human cell types. These markers were selected based on their generalizability across all contexts tested. The final senescence score was defined as a linear combination of log-transformed RNA or protein abundance weighted by their ML-derived coefficients, producing a robust, quantitative score applicable to independent datasets. This score was successfully applied to external transcriptomic and proteomic data, validating its performance and confirming its ability to accurately classify senescent cells across experimental systems. The generalizability of this ML-based scoring makes it particularly valuable for future applications in bulk and single-cell transcriptomic datasets, and in spatial profiling platforms. The independence of this ML score from cell type or stimulus-specific biases provides an orthogonal, data-driven method to complement traditional markers such as CDKN2A, CDKN1A, or LMNB1, which showed relatively low generalizability (**Figure 3**C and D).

The heterogeneity of the SASP is well documented in culture,^9^ but the heterogeneity of the intracellular proteome of senescent cells is less defined. As a result, the detection of senescence-associated proteins in tissues is largely based on more widely available RNA profiles of senescent cells originally developed in culture models, especially primary fibroblasts. However, as we report here and others have noted, transcriptomic and proteomic signatures of senescence often diverge substantially (**Figures 1**G and H; **Figure S4**D), with limited overlap in the specific markers identified in each case. This discrepancy likely reflects both technical limitations and differences between steady-state mRNA levels (the net effect of transcription and turnover) and protein levels (the net effect of translation and protein stability/processing) in senescent cells. Transcriptomic tools are, at present, more broadly applied to the spatial and single-cell omic-level mapping of senescent cells in tissues,^36^ and still require substantial improvement. In this context, proteomic approaches are critical for validating and expanding RNA-based inferences. With the advent of increasingly multiplexed, probe-based proteomic assays (GeoMX, CODEX, etc.),^37,38^ and untargeted mass spectrometry-based detection of proteins at single-cell resolution,^39,40^ wider use of proteomic methods to characterize *in vivo* senescence phenotypes, particularly the SASP, will be crucial for rigorous senotype profiling in tissues.

For a more sophisticated and robust approach to determine cell senescence, we developed an ML-guided strategy combining a minimal set of transcriptomic and proteomic features that offers a practical and scalable solution to identify senescent cells across tissues, as other labs have also begun to use, with promising outcomes. From the transcriptomes and proteomes in the rich array of senescence models represented in SenCat, we developed an ML algorithm able to identify senescent cells, both in culture and in tissues in vivo. The ML-derived marker scores outperformed canonical markers in accuracy and generalizability, successfully distinguishing senescent from non-senescent states across a range of cell types, senescence inducers, and experimental models. This approach circumvents the limitations of relying on either universal or cell type-specific markers and instead captures consistent transcriptional and proteomic patterns associated with senescence, regardless of cellular context. In this manner, it represents a powerful tool for profiling senescence in complex *in vivo* environments, where multiple senescence programs may coexist.

Nonetheless, several caveats should be considered. Importantly, senescence is a dynamic process, and the molecular features of senescent cells can vary considerably depending on their stage along the senescence trajectory; certain markers may be transient, emerging only at early or late times, and others may reflect secondary adaptations rather than core senescence. In addition, the senescence program is dependent on the nature and strength of the inducing stimulus; while our models captured generalizable signatures, they may miss markers specific to certain senescence contexts. Furthermore, the profiling method used critically influences marker selection, as we observed only partial (∼50%) overlap between transcriptomic and proteomic senescence markers, emphasizing the need to integrate multi-omic perspectives. These caveats highlight the need to refine these approaches with more temporally resolved data, a wider range of inducers, and complementary measurement platforms to enhance the accuracy and biological depth of strategies to detect senescent cells.

Recent efforts have identified transcriptomic signatures of cellular senescence derived from a different panels of human in vitro datasets.^14–17,33^ These models demonstrate impressive cross-dataset generalizability and scoring robustness, but remain constrained by a reliance on cell culture systems, a lack of *in vivo* dynamic profiling, and the absence of complementary proteomic datasets. In addition, most are trained on human cell culture data, much of which is derived from immortalized or partially transformed cell lines.^14–17,33^ Building upon this important previous work, SenCat introduces several key innovations. First, SenCat is grounded in time-resolved *in vivo* snRNA-seq datasets, enabling the identification of senescence dynamics across cell types and permitting the distinction of early changes in cell senescence from those occurring later on, both naturally and aberrantly persisting within the organs. Second, SenCat complements transcriptomic profiles with proteomic datasets from the same senescent cell types, refining our lists to those likely to include detectable secreted or surface proteins in biological fluids. Indeed, our companion publication (Olinger et al., submitted)^18^ demonstrates that the proteomic component of SenCat can be leveraged to identify clinically relevant senotype-specific signatures in human plasma proteomic data, with greater performance and tissue-resolution than previously reported. This dual-modality (proteomic and transcriptomic) design enhances the translational relevance of the catalog. Third, SenCat leverages supervised ML to derive senescence marker coefficients that are both interpretable and adaptable to specific tissues, cell types, and senescence triggers. Finally, the functional relevance of the SenCat senescence markers was supported by its ability to map ligand-receptor interactions between senescent cells and neighboring epithelial populations in old lungs. By leveraging *in vivo* snRNA-seq and curated senescence scores, SenNet identified candidate mediators of epithelial modulation by senescent cells, including potential paracrine signaling pathways. Thus, SenCat-derived ML scoring offers further insight into how senescence shapes tissue microenvironments, a feature difficult to address by existing transcriptomic models derived from cell culture systems.

Together, these features make SenCat a powerful complementary resource to existing signatures. We propose that the datasets reported here provide a robust foundation towards a consensus on the defining features and identification of senescent cells, serving as a resource to help catalyze innovation in the design of senotherapies for clinical benefit.

## METHODS

### Cell culture, senescence induction and validation

All the cultured cells in this study are of human origin. WI-38 lung fibroblasts (obtained from the NIGMS Human Genetic Cell Repository, Coriell Institute for Medical Research; repository ID AG06814-N) and BJ skin fibroblasts (ATCC, CRL-2522) were cultured in DMEM (Gibco) supplemented with 10% heat-inactivated fetal bovine serum (FBS). HSAEC lung epithelial cells (ATCC, PCS-301-010) were cultured using an Airway Epithelial Cell Basal Medium plus Bronchial Epithelial Cell Growth Kit (ATCC). HEKn epidermal skin keratinocytes (ATCC, PCS-200-010) were cultured in Dermal Cell Basal Medium plus Keratinocyte Growth Kit (ATCC). HCAEC coronary artery endothelial cells (LifeLine Cell Technology, FC-0032) were cultured in VascuLife EnGS Endothelial Medium Complete Kit (LifeLine). HUVEC umbilical vein endothelial cells (ATCC, PCS-100-010) were cultured in Vascular Cell Basal Medium plus Endothelial Cell Growth Kit-BBE (ATCC). BMMSC bone marrow mesenchymal cells (ATCC, PCS-500-012) were cultured in Mesenchymal Stem Cell Basal Medium for Adipose, Umbilical and Bone Marrow-derived MSCs plus Mesenchymal Stem Cell Growth Kit for Bone Marrow-derived MSCs (ATCC). HVSMC coronary artery smooth muscle cells (LifeLine Cell Technology, FC-0031) were cultured in VascuLife SMC Medium Complete Kit (LifeLine). HSKM skeletal myoblasts (Gibco, A12555) were cultured in an equal volume mixture of Ham’s F10 medium supplemented with 20% fetal bovine serum (FBS) and Promocell skeletal muscle cell growth medium. PBMC peripheral blood mononuclear cells (ATCC, PCS-800-011) were cultured in RPMI 1640 plus 10% FBS. PreAdipo subcutaneous preadipocytes (ATCC, PCS-210-010) were cultured in Fibroblast Basal Medium plus Fibroblast Growth Kit-Low Serum (ATCC). NHO osteoblasts (Lonza, CC-2538) were cultured in OGM Osteoblast Growth Medium BulletKit (Lonza). NHA astrocytes (Lonza, CC-2565) were cultured in ABMTM Basal Medium plus AGMTM SingleQuotsTM Supplements (Lonza). HEMn melanocytes (ATCC, PCS-200-012) were cultured in Dermal Cell Basal Medium plus Melanocyte Growth Kit (ATCC).

Culture media were supplemented with 0.5% penicillin-streptomycin (25 U/mL penicillin and 25 µg/mL streptomycin final concentrations), sodium pyruvate, and non-essential amino acids (Gibco), in a 5% CO_2_ incubator. At the time of inducing senescence, all cells were at a low passage and ∼50% confluence. Further details on senescence triggers, doses, and duration of the treatments are in **Table S9**. Specifically, PBMCs were cultured for 6 days at a concentration of 1.4 million cells/mL of medium for senescence induction. The senescent state was confirmed by measuring BrdU incorporation to assess proliferation (Cell Signaling Technology, 6813) and increased SA-β-gal activity (Cell Signaling Technology, 9860) following the manufacturer instructions. A ‘cell type marker’ refers to those markers shared by all senescence models characterized in that cell type. A ‘senescence model’ is a particular cell type in which senescence was triggered with a specific treatment. To minimize batch effects, we collected and processed experimental samples from each cell line at the same time.

### Determination of transcriptomic and proteomic markers (no ML)

Every transcript or protein that significantly (p-value < 0.05) increased or decreased by at least 1.5-fold (Log2 FC 0.585) was included in the marker lists for every senescence model. Senescence markers for a particular cell type were determined by selecting only those present in all the lists for every senescence model (every trigger used). From the transcriptomic analysis, normalized counts higher than 1000 in the target population (>1000 in CTIS for the transcripts up, <1000 in P/EV for the transcripts down) were included. The top 20 shared senescence markers were not ranked by FC; given that no markers were shared by all cell types, these lists were ranked by the number of cell types in which these markers were present. On the other hand, each list of the top 20 cell type-specific senescence markers was ranked by the sum of their Log2FC values. Expanded lists of cell type-specific senescence markers used for scRNA-seq analysis considered all the models separately from each cell type (e.g., ETIS, IRIS, OSIS, and OIS from both WI-38 and BJ separately). The score lists were made including as many genes as possible given the nature of the calculating function; the more RNAs to consider the better, as it is an average expression function (AddModuleScore in Seurat). For the lists of mouse RNAs, homologous transcripts were obtained from the NIH’s Gene platform (ncbi.nlm.nih.gov/datasets/gene), including all available ortholog mappings (not only 1:1 relationships but also 1:many and many:many) to retain maximal biological information when translating human marker sets to mouse genes. Given the nature of the algorithm evaluating the expression score in samples profiled by scRNA-seq analysis, which calculates the average levels of all of the RNAs in the list, we considered that a longer list would more reliably detect a “senescent-like” cluster. Therefore, we created expanded lists of the transcripts present in at least 4 senescent cell types (“Allsenshared”) and the top 500 present in more senescence models per cell type (“Expandedtopsenfib” for fibroblasts, “Expandedtopsenendo” for endothelial cells, and “Expandedtopsenepi” for epithelial cells).

### Machine learning (ML) determination of senescence markers

The SenCat transcriptomic and proteomic datasets were used for the purpose of obtaining ML-based lists of senescence markers applicable to all models. For transcriptomic analyses, Ensembl gene identifiers were used, and only features (RNAs mapped to genes) with a single ID were retained. Proteomic data were annotated using UniProt identifiers.

For each omics layer, features were filtered to retain only those (1) with non-zero expression values across all samples and (2) with expression above the 80^th^ percentile in the full dataset. Remaining features were log1p-transformed. From these, the top 1000 features were selected using SelectKBest from the sklearn.feature_selection module with default parameters. We implemented a leave-one-cell-type-out strategy, training 14 independent binary classifiers to distinguish senescent from non-senescent samples. Each classifier excluded one cell type from training and used it for testing. Models were trained using LogisticRegression from sklearn.linear_model with L1 regularization (penalty=’l1’, solver=’liblinear’).

From each model, features with absolute weight >0.05 were extracted. Only features consistently retained as informative across all 14 models were considered robust markers and included in the final RNA/protein set. To generate interpretable, generalizable marker weights, the analysis was repeated using only the intersection of features selected by all models. Next, 14 new logistic regression models were trained (one per leave-one-out split), this time disabling regularization (penalty=None) to allow all retained markers to be assigned a non-zero coefficient. Final weights were computed as the mean coefficient across all models for each marker.

The final transcriptomic and proteomic scores were validated in an external dataset of IMR-90 lung fibroblasts, which included proliferating (P), quiescent (QUI), and senescent (SEN) conditions. We applied the same normalization and scoring strategy to this dataset without retraining the model.

### Principal component and UMAP analyses of bulk transcriptomic and proteomic data

To assess global patterns in transcriptomic and proteomic senescence profiles, we performed principal component analysis (PCA) and uniform manifold approximation and projection (UMAP) analysis on bulk RNA-seq and proteomics datasets.

Transcriptomic data were obtained from **Table S1** and proteomic expression data from **Table S2**. Before dimensionality reduction, each dataset was log2-transformed and scaled. For PCA, the top 2,000 most variable transcripts and the top 11,000 most variable proteins were selected based on their variance across samples, as determined by elbow plots (variance-ranked feature curves). PCA was performed using the prcomp() function in R, with centering and scaling applied. UMAP analysis was performed using the umap() function from the uwot R package and establishing settings provided an interpretable balance between local and global structure preservation. Metadata (cell type and treatment) were extracted from sample identifiers and used to annotate UMAP and PCA projections. Full analysis scripts and feature selection criteria are available upon request. All dimensionality reduction analyses were performed independently for transcriptomic and proteomic datasets to capture modality-specific structure.

### Bulk RNA-seq analysis

Bulk RNA was extracted using RLT buffer (Qiagen) on the QIAcube system (Qiagen), following the RNeasy Plus protocol. RNA quality and quantity were assessed with Agilent RNA Screen Tape on the Agilent Tapestation. High-quality RNA (150 ng) was used for library preparation following the Illumina TruSeq Stranded Total RNA Library Preparation Kit protocol (Illumina, Cat# 20020598). Briefly, after rRNA depletion and cDNA generation, the cDNAs were subjected to 3’-end adenylation, adapter ligation, purification with AMPure beads (Beckman, Cat#A63881), and size selection of the products with SPRIselect beads (Beckman, Cat#B23318). The selected cDNAs were enriched by PCR amplification and purified again with SPRIselect beads to generate sequencing libraries. The quality and quantity of the sequencing libraries were checked using Agilent DNA 1000 Screen Tape on the Agilent Tapestation. Paired-end sequencing was performed for ∼110 cycles on Illumina Novaseq 6000 sequencer with a depth of ∼200 million reads per sample. Following RNA-sequencing, the BCL files were demultiplexed and converted to FASTQ files using the bcl2fastq program (v2.20.0.422). The quality of the sequence data in FASTQ files was evaluated with FastQC (v0.11.9), and the reads were trimmed using BBDuk from BBTools (v38.76) and mapped to the human reference genome hg38 (Ensembl v104) using STAR aligner (v2.7.0f_0328).^41^ The reads were quantified with featureCounts in R package Subread (v2.10.5),^42^ and the resulting counts were normalized and used for differential expression analysis between experimental groups with the DESeq2 package (v1.32.0).^43^ Analyses shown in **Figure 3** and **S6** were carried out by using each cell type’s senescence markers (**Figure 3**A, 3D and **S6**A,D) and all the senescence markers shared by at least 4 cell types (**Figure 3**B, 3C, 3E, 3F and **S6**B, S6C, S6E, S6F) on EnrichR^44^ and Metascape^45^ platforms.

### Single-nuclei (sn)RNA-seq analysis

Nuclei were isolated from frozen mouse tissues using a combination of mechanical homogenization and the Minute™ Single Nucleus Isolation Kit for Tissues/Cells (Invent Biotechnologies, SN-047).

For kidney samples, six mouse kidneys were minced on dry ice using a sterile scalpel in a 10-cm Petri dish. The tissue was homogenized in 5 mL of ice-cold homogenization buffer (10 mM Tris-HCl, 10 mM NaCl, 3 mM MgCl_2_, 0.1% Nonidet™ P40 Substitute [Sigma-Aldrich, #74385] in nuclease-free water) until no visible chunks remained. Homogenates from three mice were combined to generate a pooled sample. For lung tissue, approximately 50 mg of frozen tissue was processed per sample using the same homogenization procedure.

A 200-µL aliquot of the pooled homogenate was transferred to a 1.5 mL DNA LoBind tube (Eppendorf, #022431021), mixed with 400 µL of cold lysis buffer from the Minute™ kit, and incubated on ice for 10 min. Following incubation, the lysate was passed through a proprietary filtration column and centrifuged at 600 × g for 5 min at 4°C. The supernatant was discarded, and the pellet was resuspended in 0.5 to 0.8 mL of cold wash buffer (1× PBS, 1.0% BSA [Ambion, AM2616], 0.2 U/μL RNase Inhibitor [Sigma-Aldrich, #3335399001]) by repeated pipetting. After centrifugation at 500 × g for 5 min at 4°C, the nuclei pellet was resuspended in 50-200 µL of PBS with 1% BSA and 0.2 U/µL RNase Inhibitor. Nuclei concentration was determined by mixing the sample 1:1 with Trypan Blue Stain (0.4%) and counting using a hemocytometer. Nuclei were diluted to a final concentration of 700-1200 nuclei/µL and immediately processed for snRNA-seq analysis.

Nuclei were loaded into a Chromium™ Controller (10x Genomics) for partitioning into Gel Bead-in-Emulsions (GEMs) using the Chromium GEM-X Single Cell 3’ Kit v4 (PN-1000691). Reverse transcription and barcoding of individual nuclei were performed within GEMs, followed by library construction using the Chromium Single Cell 3’ Library Construction Kit C (10x Genomics, PN-1000694), following the manufacturer’s protocols. Final libraries were sequenced on an Illumina NovaSeq X-Plus platform using paired-end 150-bp reads, with a targeted sequencing depth of approximately 20,000 reads per nucleus.

Sequencing data were then processed using Cell Ranger v9.0.0 (10x Genomics) with the --include-introns option enabled to support nuclear transcript detection. Reads were aligned to the mouse reference transcriptome GRCm39-2024-A, and gene expression matrices were generated using default parameters. Key quality control metrics, such as mapping rate, number of detected genes per nucleus, and sequencing saturation, were obtained from Cell Ranger web summary reports.

The resulting matrix files were processed in R (v4.5.0) using Seurat (v5). From the lung time course samples, all “Day 12” replicates along with 1 replicate of the “Day 1” condition were removed from the analysis due to technical issues. Cells with >200 and <10000 nFeature_RNA, <25000 nCount_RNA, and <25% mitochondrial genes were included for analysis. Doublets were removed by using scDblFinder function. Whole samples were merged and normalized, and the top 2000 highly variable RNAs were retained. Data from each experiment were scaled, dimensionally reduced by PCA, and integrated through the Harmony package. Clustering was then performed and UMAP plots prepared; 21 clusters were identified per tissue and experiment. Cell types were identified by curating the markers obtained through the FindAllMarkers function with GPT-4, and curated with markers identified in previous works from our group. Sequencing data are available at GSE302792 (token szybiqiozdcpjel).

To identify senescent cells in lung and kidney, we first applied a multi-step strategy combining marker refinement, machine learning, and UMI normalization to score senescence at single-cell resolution. Starting from the full ML-identified list of senescence-associated genes (SenCat ML all), we removed potential false positives by excluding genes aberrantly upregulated in control sample cells within any cluster (minimal senescence conditions, Day 0 in samples from doxorubicin time course, young mice in natural aging samples). Specifically, we performed cluster-wise differential expression analyses between Day 0 cells and all other time points, retaining only genes that were not significantly upregulated in Day 0 in any cluster. We then trained a LASSO-regularized logistic regression model using log-normalized expression values of the filtered genes across all cells. To emphasize the specificity of senescence, control sample cells were labeled as non-senescent (0), while all others were labeled as senescent (1). A small subset of the control sample cells was randomly relabeled as “senescent” (1) to avoid complete separation and improve model robustness. We applied sample weights to downweigh control sample cells and used 10-fold cross-validation to determine the optimal regularization parameter. The resulting model coefficients defined a refined senescence gene set-based on the non-zero weighted predictors.

This refined gene set was then used to compute cell-wise enrichment scores using the AUCell algorithm, which ranks transcriptomes within each cell and calculates the area under the recovery curve for the input transcriptome set. To correct for technical confounding due to sequencing depth, we performed a *post hoc* correction of the AUCell scores using a generalized additive model (GAM) with a spline term for nCount_RNA (UMI counts). The residuals from the GAM were stored as corrected senescence scores. We used only markers that increased in the snRNA-seq analysis, as this approach is not yet adequate to identify reduced markers, and it may therefore introduce noise. Moreover, AUCell is not able to compute scores with reduced RNAs.

Finally, to define senescent cells, we established a threshold based on the 99^th^ percentile of corrected AUCell scores from control sample cells. Cells exceeding this threshold were labeled as senescent. The proportion of senescent cells was then calculated across conditions and clusters to characterize the distribution and dynamics of senescence across the different samples included in the analyses. We then extracted the differential expression (log-normalized FC) of the cells identified as senescent versus non-senescent per cluster, and those data were analyzed through GSEA for pathway and gene set comparison between conditions.

### Sample preparation for mass spectrometry

Cell pellets were processed for downstream mass spectrometry (MS)-based proteomic analysis by a standard in-solution digestion protocol, as previously described.^46^ Briefly, cell pellets were solubilized in a lysis buffer composed of 8 M Urea in 50 mM triethylammonium bicarbonate (ThermoFisher #90114). For each sample, 50 to 100 µg of protein, based on BCA analysis (ThermoFisher #23225), was processed by in-solution digestion. Protein samples were incubated in 20 mM dithiothreitol (Sigma #D9779) for 30 min at 37°C to reduce disulfide bonds, followed by incubation in 40 mM iodoacetamide (Sigma #I1149) for 30 min at room temperature in the dark to irreversibly alkylate sulfhydryl groups. Samples were then diluted below 1 M urea with 50 mM triethylammonium bicarbonate and digested into peptides with trypsin (Promega #V5113) at a 1:50 trypsin:protein ratio, by mass, overnight at 37°C with agitation. Digestion was quenched with 1% formic acid (Sigma #27001). Peptide samples were desalted with Oasis HLB Solid Phase Extraction Cartridges (Waters #186000383) according to the manufacturer protocol. Samples were then dried in a vacuum concentrator and resuspended in mass spectrometry sample loading buffer (0.2% formic acid in water).

### Liquid chromatography-mass spectrometry

All samples were analyzed using an UltiMate 3000 nano-HPLC system coupled to a hybrid Orbitrap Eclipse mass spectrometer (ThermoFisher) with an EASY-Spray source (Thermo Fisher Scientific), as previously described.^47^ Digested peptides (1 µg of each sample) were loaded on a trap column (75 μm × 20 mm, 3 μm C18 particle) at a constant flow rate of 5 μL/min and separated on a nano column (75 μm × 500 mm, 2 μm C18 particle, ThermoFisher #ES903) using a 2-h efficient linear gradient with constant flow rate of 300 nL/min, 0-5 min (2% phase B, sample loading), 5-120 min (2%-35% phase B), 120-125 min (35%-80% phase B), 125-135 min (80% phase B), 135-136 min (80%-2% phase B), 136-150 min (2% phase B), phase A contains 5% DMSO in 0.1% formic acid water, and phase B contains 5% DMSO in 0.1% formic acid ACN. The nano and trap columns were heated to 60°C in the EASY-Spray electronic ionization source and column oven, respectively.

The MS data were acquired in data-independent acquisition (DIA) mode. MS1 spectra were acquired in the Orbitrap at 240,000 resolution over 350-1000 m/z range. The MS1 normalized AGC target was set to Standard with Max Injection Time set to Auto. For MS/MS scans, the precursor range was set to 400-1000 m/z, and DIA experiment was performed with 8 m/z isolation windows stepped across the precursor range with a 1 m/z overlap, resulting in 75 windows for each scan cycle. MS2 scans were collected from 150-2000 m/z. MS2 fragmentation was performed using HCD with 30% collision energy. DIA MS2 scans were acquired in Orbitrap analyzer at 30K resolution with a normalized AGC target of 800%. The MS2 scan range was defined as 145-1450 m/z, and loop control was set to 3 s. For MS acquisition, samples were run in a block-randomized design to minimize batch effects.

### Protein identification and quantitative analysis pipeline

Raw DIA data files were analyzed using the Spectronaut software (v18.1) utilizing the directDIA algorithm with default settings. The Uniprot human fasta database (UP000005640_20191105) was used. Carbamidomethylation was set as a fixed modification. Methionine oxidation and N-terminal acetylation were set as variable modifications. FDR was set to 1% for filtering identifications at the peptide and protein group levels. All settings were otherwise set to default. Differential analysis was conducted on the protein intensities between senescent and non-senescent samples using the ProtPipe package in R,^48^ using peak area-based protein-level quantitation reports generated by Spectronaut as input data. Multiple testing correction was performed using the Benjamini-Hochberg procedure. Raw proteomic data are available from MassIVE (ftp://massive.ucsd.edu/v07/MSV000096215/); dataset identifier MSV000096215.

### RT-qPCR analysis

Cells were lysed in RLT buffer (Qiagen). Lysates were processed with the QIAcube (Qiagen) to purify total RNA, and then reverse transcribed to synthesize cDNA using Maxima reverse transcriptase (Thermo Fisher Scientific) and random hexamers. Real-time, qPCR analysis was performed using SYBR Green mix (Kapa Biosystems), and the relative mRNA levels were determined by the 2^−ΔΔCt^ method. All mRNAs evaluated were normalized to human *ACTB* mRNA or mouse *Actb* mRNA levels. Primers used to detect human mRNAs were as follows (each pair forward and reverse, respectively): GCACAGAGCCTCGCCTT and GTTGTCGACGACGAGCG for *ACTB* mRNA; GTTACGGTCGGAGGCCG, and GTGAGAGTGGCGGGGTC for *p16/CDKN2A* mRNA; AGTCAGTTCCTTGTGGAGCC and CATGGGTTCTGACGGACAT for *CDKN1A* mRNA; AGTGAGGAACAAGCCAGAGC and GTCAGGGGTGGTTATTGCAT for *IL6* mRNA; GACCCTCAGAGTTGCACTCC and GCCTGGTTAGCAGGTCCTC for *GDF15* mRNA; and GAAAAAGACAACTCTCGTCGCA and GTAAGCACTGATTTCCATGTCCA for *LMNB1* mRNA.

### Mice and histology

All mouse procedures, including import, housing, experimental work, and euthanasia, were conducted under Animal Study Proposal # 476-LGG-2027, reviewed and approved by the Animal Care and Use Committee of the NIA, NIH. C57BL/6JN mice were provided the Inotiv 2018SX diet *ad libitum* and kept on a 12 h-12 h light-dark cycle. All drugs used in the study were administered intraperitoneally with a maximum volume of 80 µL. Doxorubicin-induced senescence was triggered by a single dose of doxorubicin (10 mg/kg, Selleckchem) in 3-month-old mice and studied 8 days later. DMSO or ABT-737 (25 mg/kg, MedChemExpress) were administered to mice treated with doxorubicin according to specified regimens.

## ACKNOWLEDGEMENTS

This work was supported by the National Institute on Aging (NIA) Intramural Research Program (IRP), NIH. RdC and NB were supported by a SenNet NIH Common Fund Grant (NIA U54 AG079779, PI: Elisseeff). CA was supported by a fellowship from the NIA IRP, NIH, a “Cesar Nombela” fellowship from the autonomous government of Madrid (2023-T1/SAL-GL-29093), a fellowship from the “la Caixa” Foundation (ID 100010434, LCF/BQ/PI24/12040003), and by the European Union (ERC, ChECMate senescence, 101163448).

## AUTHOR CONTRIBUTIONS

CA and AH conceived the study. CA designed and coordinated the study. CA, MG, AH, NB, YAQ, and KA interpreted the results. CA, DT, JLM, KA, RdC, ABH, NB, and MG contributed conceptually. CA, DT, RB, GA, MSM, JHY, CYC, RM, MR, ABH, and NB developed senescence models. Transcriptomic and proteomic analysis was carried out by CA, DT, RB, ASGC, BO, QS, DVR, YH, ZL, YAQ, YP, JF, SD, and NB. CA, ASGC, and YAQ validated the findings. KG and MM performed the ML analysis. CA and MG wrote the manuscript.

