## Supplementary material for "SenCat: Cataloging human cell senescence through multiomic profiling of multiple senescent primary cell types": Legend to Supplemental Figures

**SUPPLEMENTARY FIGURE LEGENDS**

**Figure S1. BrdU incorporation analysis in the SenCat senescence paradigms**

Heatmap representing the row Z-score values of the BrdU incorporation-dependent absorbance for each of the samples assessed in the SenCat project.

**Figure S2. SA-β-Gal staining to validate cell senescence in the SenCat senescence paradigms**

Representative SA-β-Gal staining images of the different conditions profiled in the SenCat project. Scale bar, 100 µm.

**Figure S3. Assessment of senescence canonical RNA markers by RT-qPCR analysis**

The levels of the indicated transcripts (canonical senescence RNA markers ) were measured by RT-qPCR analysis in the indicated conditions. All the graphs displayed in this figure show each individual value as a dot and the means ±S.D.; significance (*P < 0.05, **P < 0.01, ***P < 0.001) between each senescence model and their respective controls was calculated using a two-tailed Student’s t-test.

**Figure S4. Principal component analysis (PCA) of transcriptomic data from senescence models**

(A) PCA plots of transcriptomic profiles for each of the 14 primary human cell types included in the SenCat dataset, colored by treatment condition. Each panel displays samples from one cell type treated with at least two senescence inducers (CTIS, IRIS, OSIS, OIS) and their respective controls (P = proliferating, EV = empty vector).

(B) PCA of the entire transcriptomic dataset across all cell types and treatment conditions. Samples are colored by cell type and shaped by treatment.

**Figure S5. Principal component analysis (PCA) of proteomic data from senescence models**

(A) PCA plots of proteomic profiles for each of the 14 primary human cell types included in the SenCat dataset, colored by treatment condition. Each panel displays samples from one cell type treated with at least two senescence inducers (CTIS, IRIS, OSIS, OIS) and their respective controls (P = proliferating, EV = empty vector).

(B) PCA of the entire proteomic dataset across all cell types and treatment conditions. Samples are colored by cell type and shaped by treatment.

**Figure S6. Overlap of transcriptomic and proteomic senescence markers across triggers, cell types, and profiling approaches**

(A–D) Venn diagrams illustrating the overlap of upregulated (A, B) and downregulated (C, D) markers across three senescence triggers (CTIS, IRIS, OSIS) within each cell type, separately for transcriptomic (A, C) and proteomic (B, D) datasets. Percentages indicate the proportion of shared markers relative to the total for each cell type. Each circle represents one trigger, and overlaps indicate shared markers between conditions.

(E–F) Venn diagrams showing the number and percentage of markers shared by at least four cell types across all senescence triggers, separated by transcriptomics and proteomics for upregulated (E) and downregulated (F) markers. Marker sharing across individual cell types is visualized next to each diagram.

(G) Barplot comparing the number of shared markers between transcriptomic and proteomic datasets for upregulated and downregulated genes. It shows each individual value as a dot; the means ±S.D., significance (ns, not significant) was calculated using a two-tailed Student’s t-test.

**Figure S7. Extended data on newly discovered downregulated senescence markers and the individual performance of the upregulated markers versus the canonical ones**

(A-B) Heatmaps showing log2 fold changes of the top 20 newly-discovered senescence markers across senescence models for transcriptomic (A) and proteomic (B) analyses. Columns represent individual markers; rows indicate each cell type and senescence trigger. Grey cross marks in (A) denote conditions where RNA for the corresponding gene was not detected. Red indicates higher expression in senescent cells versus controls, while blue indicates lower expression. (C,D) Log2 FC values of each of the indicated top 20 markers (New) and the canonical ones (Canonical, highlighted by a grey box) for each senescence model (C, Transcriptomic; D, Proteomic).

**Figure S8. Pathway analysis of reduced markers of senescence**

(A) Heatmap displaying the association score (-Log10 p-value) of each cell type’s downregulated transcriptomic senescence markers with the indicated “Hallmark” pathways.

(B) Dot plots (top) and marker-pathway association grids (bottom) were created using EnrichR, and show the association between the downregulated transcriptomic senescence markers shared by at least 4 cell types with the indicated “Hallmark” and “GO Cellular Component” pathways (top dot plots). Bottom grids display 50 markers associated with a specific pathway from those categories.

(C) Metascape plot (including pathway categories only in the analysis) showing each of the pathways associated with the downregulated transcriptomic senescence markers shared by at least 4 cell types, clustered by common pathways and broader processes.

(D-F) D-F are the same as A-C, but the analysis was carried out using proteomic data instead.

**Figure S9. Extended data on the transcriptomic ML strategy in cultured cells**

Heatmaps of traditional senescence markers (p16, p21, IL6, GDF15, LMNB1) and ML-derived transcriptomic markers (TOP2A, MX1, GDF15, CXCL8, H1-5, H3C2, RN7SL471P, H3C14, H3C15) and senescence scores (ML T SENCAT) across 14 different human cell types subjected to senescence-inducing treatments (CTIS, IRIS, OSIS, OIS) or controls (P or EV) from the transcriptomic data.

**Figure S10. Extended data on the proteomic ML strategy in cultured cells**

Heatmaps of traditional senescence markers (p16, p21, IL6, GDF15, LMNB1) and ML-derived transcriptomic markers (FDXR, SERPINH1, STMN1, VDAC2, H1-5, TAGLN, SQOR, PCNA, ATP5MG, H4C1, GAA, PHGDH) and senescence scores (ML P SENCAT) across 14 different human cell types subjected to senescence-inducing treatments (CTIS, IRIS, OSIS, OIS) or controls (P or EV) from the transcriptomic data.

**Figure S11. Senescence scoring and signature comparison in lung samples**

(A) UMAP projections of lung samples across time points, colored by cluster.

(B) Density plots of AUCell scores using various senescence signatures across time points.

(C) Table summarizing total and senescent cell counts and percentages across conditions.

(D) UMAP projections showing expression of canonical senescence markers across time points.

**Figure S12. Application of the senescence scoring strategy to doxorubicin-treated kidney time course samples**

(A) UMAP projection of kidney samples colored by cluster.

(B) Table of cluster identities in kidney samples.

(C) UMAP projections of kidney samples across time points.

(D) UMAP projections showing expression of canonical senescence markers across time points.

(E) Density plot of AUCell scores using the refined “SenCat ML all” signature.

(F) UMAP projections showing distribution of senescent cells based on refined “SenCat ML all” scores.

(G) Table summarizing total and senescent cell counts and percentages across conditions.

(H) Barplot showing the number of senescent cells per cluster and condition in kidney samples.

**Figure S13. Senescence scoring and cell-cell interaction analysis in lung from old mice**

(A) UMAP projections of lung samples from 3-, 12-, and 23-month-old (mo) female mice.

(B) Table of cluster identities in lung samples from old mice.

(C) UMAP projections of lung samples from old mice, colored by cluster.

(D) Density plot of AUCell scores using the refined “SenCat ML all” signature across age groups.

(E) Table summarizing total and senescent cell counts and percentages per age group.

(F) Barplot showing senescent cells per cluster and aging group.

(G) Table listing inferred ligand–receptor interactions between senescent fibroblasts and epithelial cells.
