## Supplemental Figures for "SenCat: Cataloging human cell senescence through multiomic profiling of multiple senescent primary cell types"

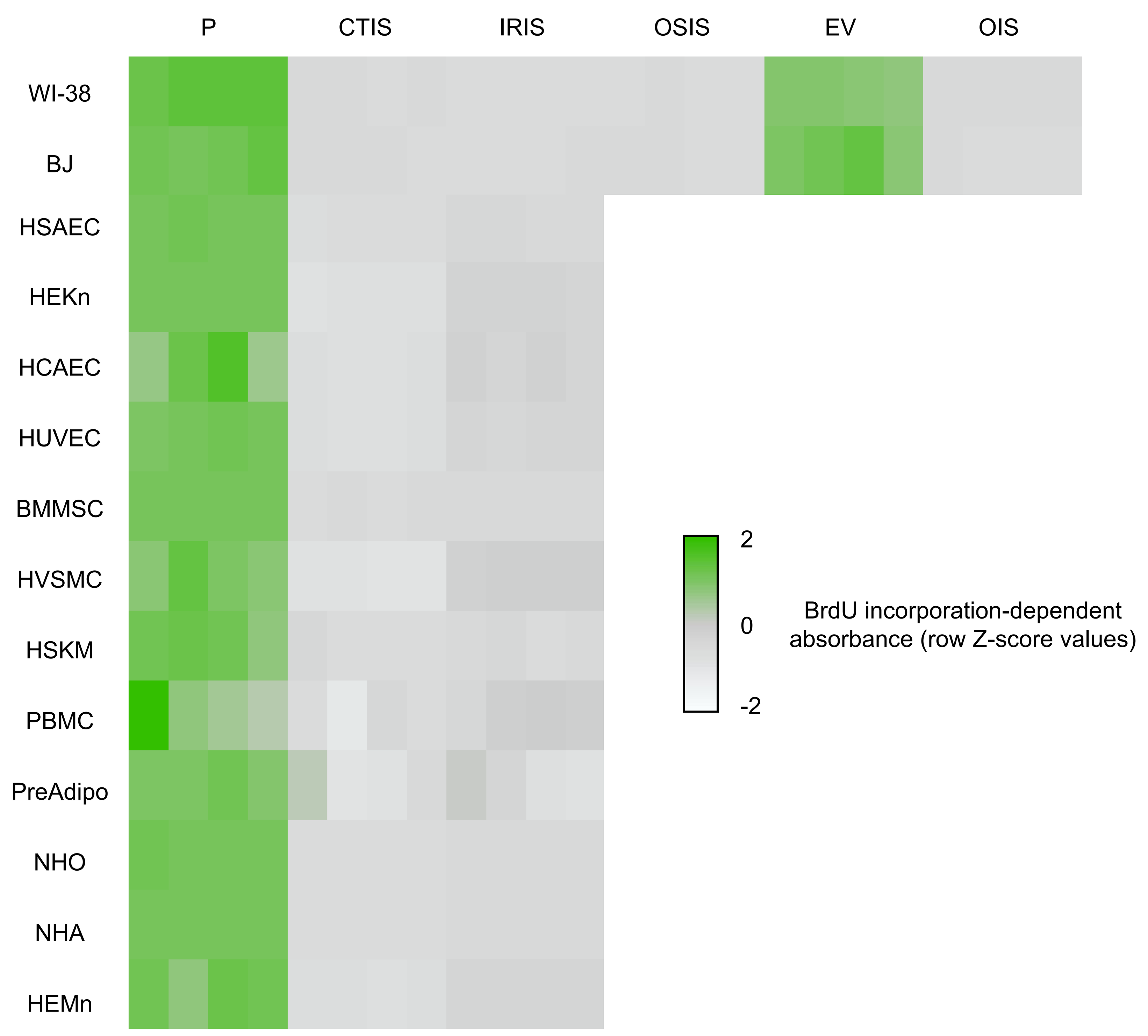
P
CTIS
IRIS
OSIS
EV
OIS
WI-38
BJ
HSAEC
HEKn
HCAEC
HUVEC
BMMSC
HVSMC
HSKM
PBMC
PreAdipo
NHO
NHA
HEMn
2
0
-2
BrdU incorporation-dependent
absorbance (row Z-score values)

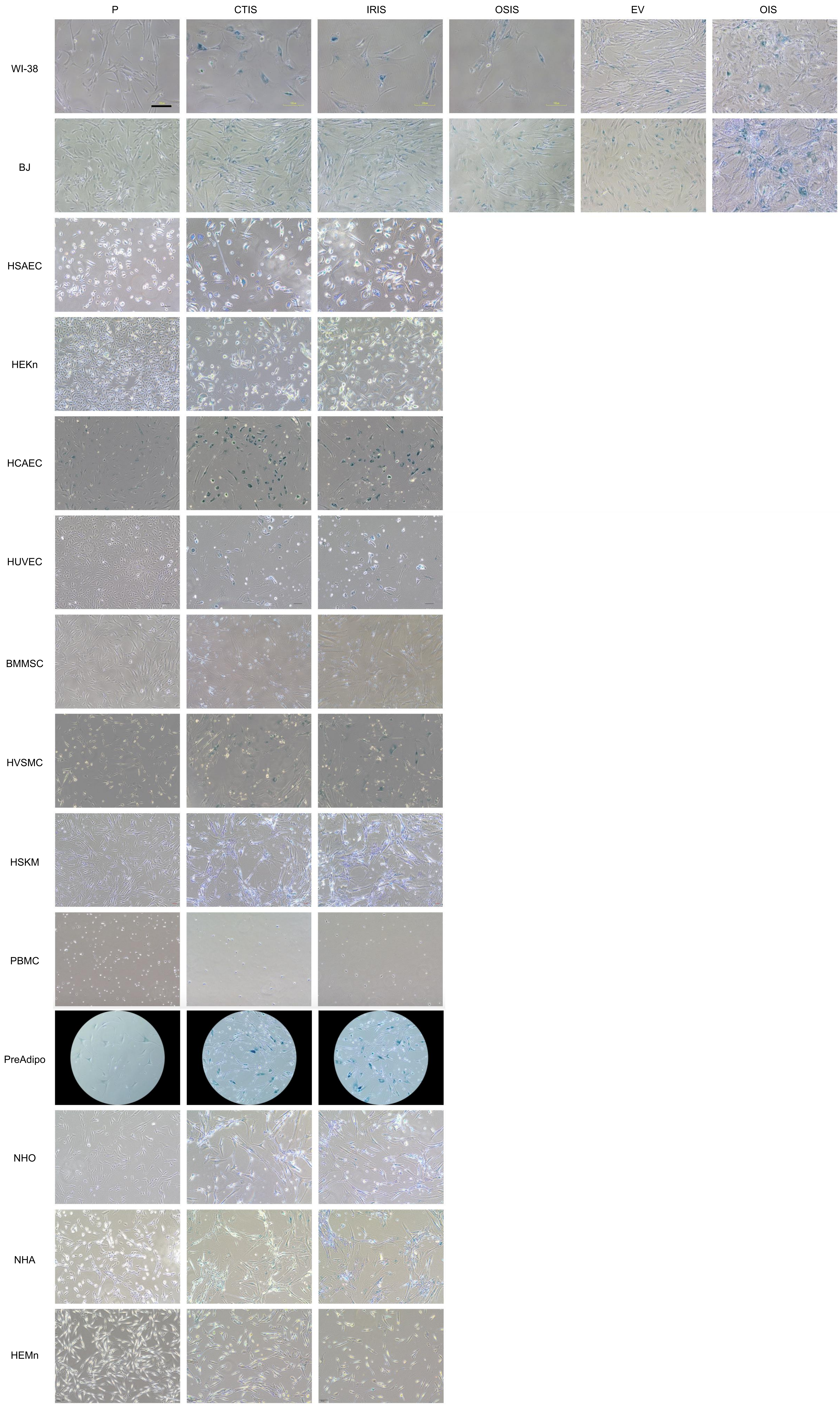
P
CTIS
IRIS
OSIS
EV
OIS
WI-38
BJ
HSAEC
HEKn
HCAEC
HUVEC
BMMSC
HVSMC
HSKM
PBMC
PreAdipo
NHO
NHA
HEMn

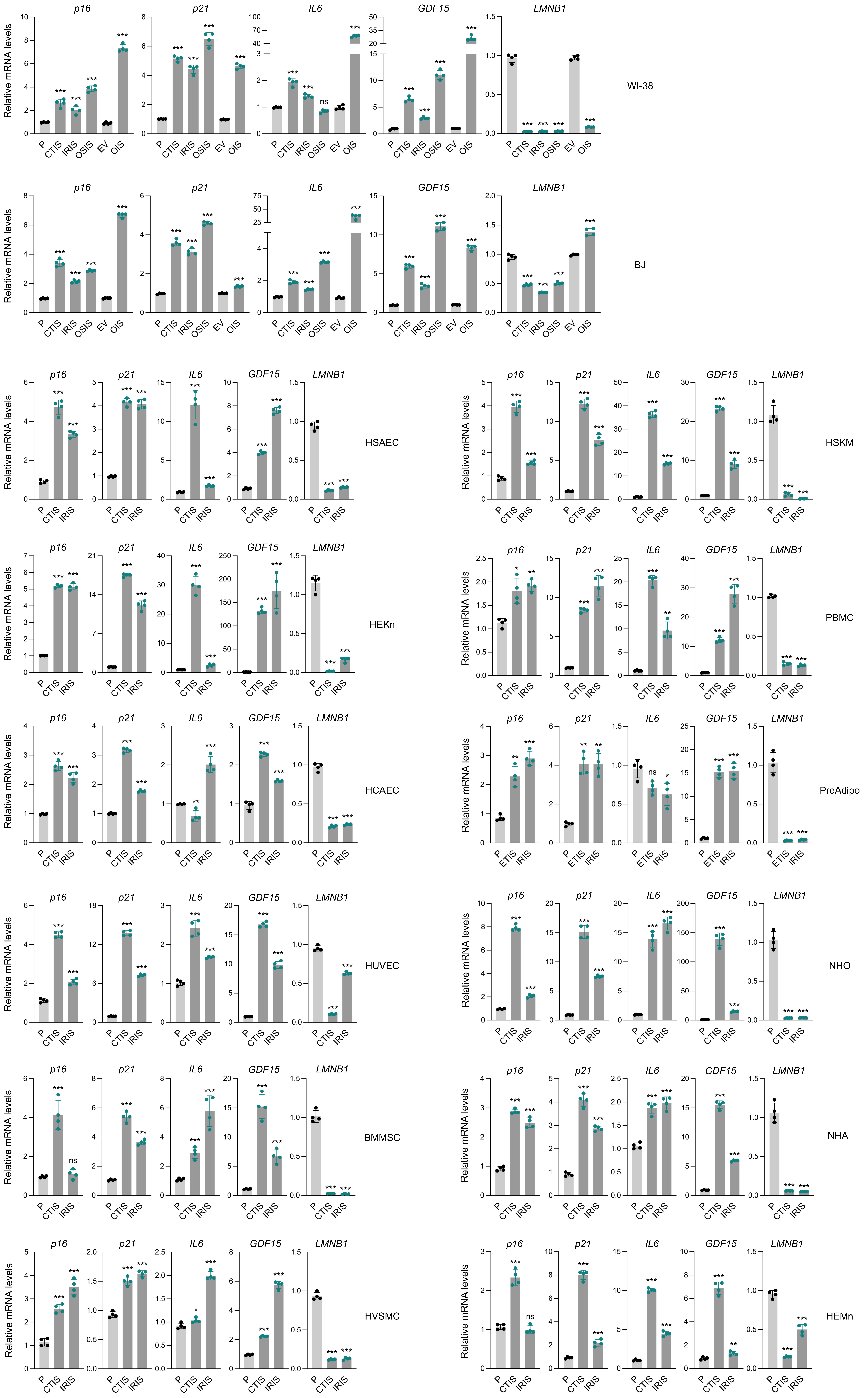
p16
p21
IL6
GDF15
LMNB1
Relative mRNA levels
P
CTIS
IRIS
OSIS
EV
OIS
ETIS
WI-38
BJ
HSAEC
HSKM
HEKn
PBMC
HCAEC
PreAdipo
HUVEC
NHO
BMMSC
NHA
HVSMC
HEMn

**A**

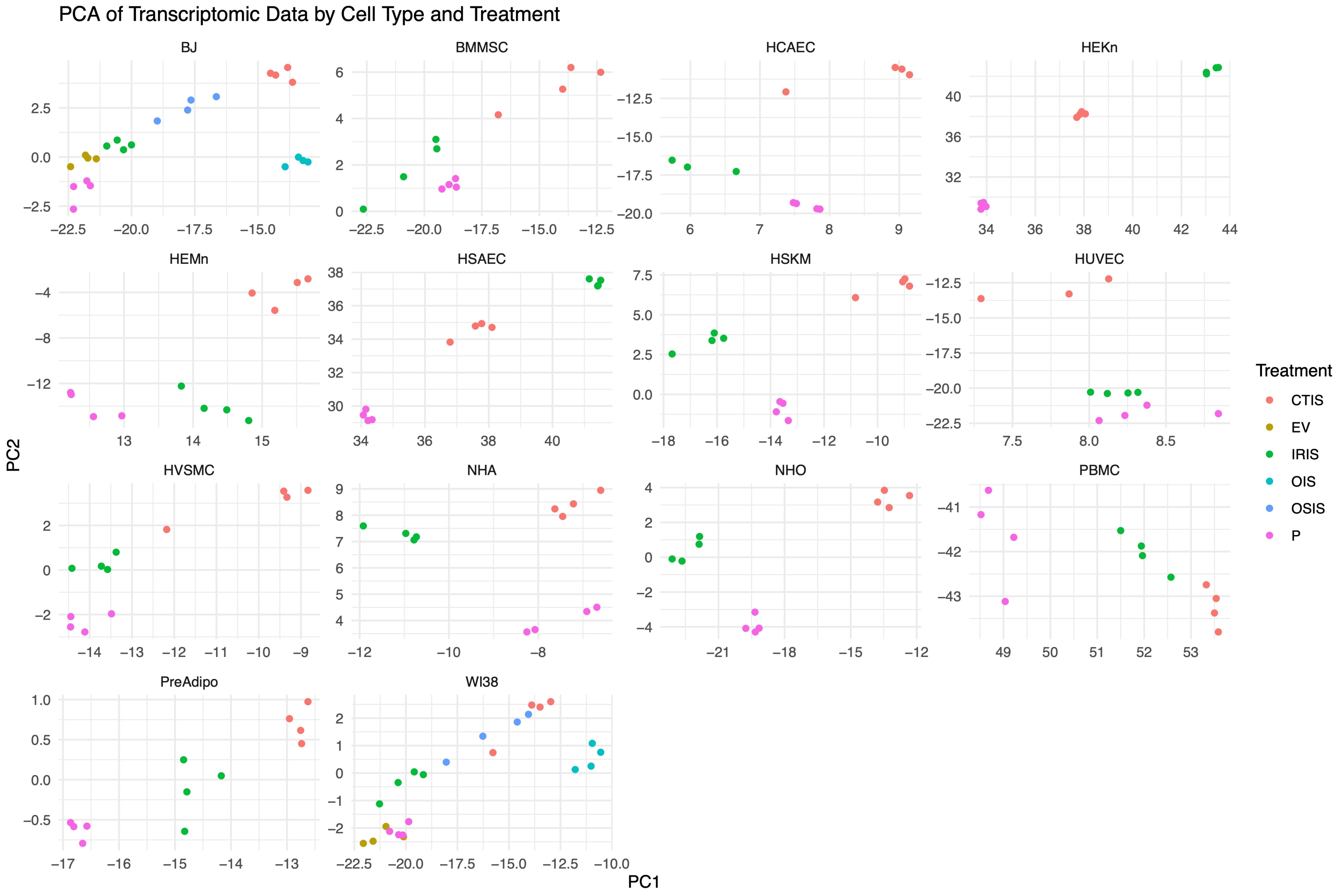
PCA of Transcriptomic Data by Cell Type and Treatment
BJ
BMMSC
HCAEC
HEKn
HEMn
HSAEC
HSKM
HUVEC
HVSMC
NHA
NHO
PBMC
PreAdipo
WI38
PC1
PC2
Treatment
CTIS
EV
IRIS
OIS
OSIS
P

**B**

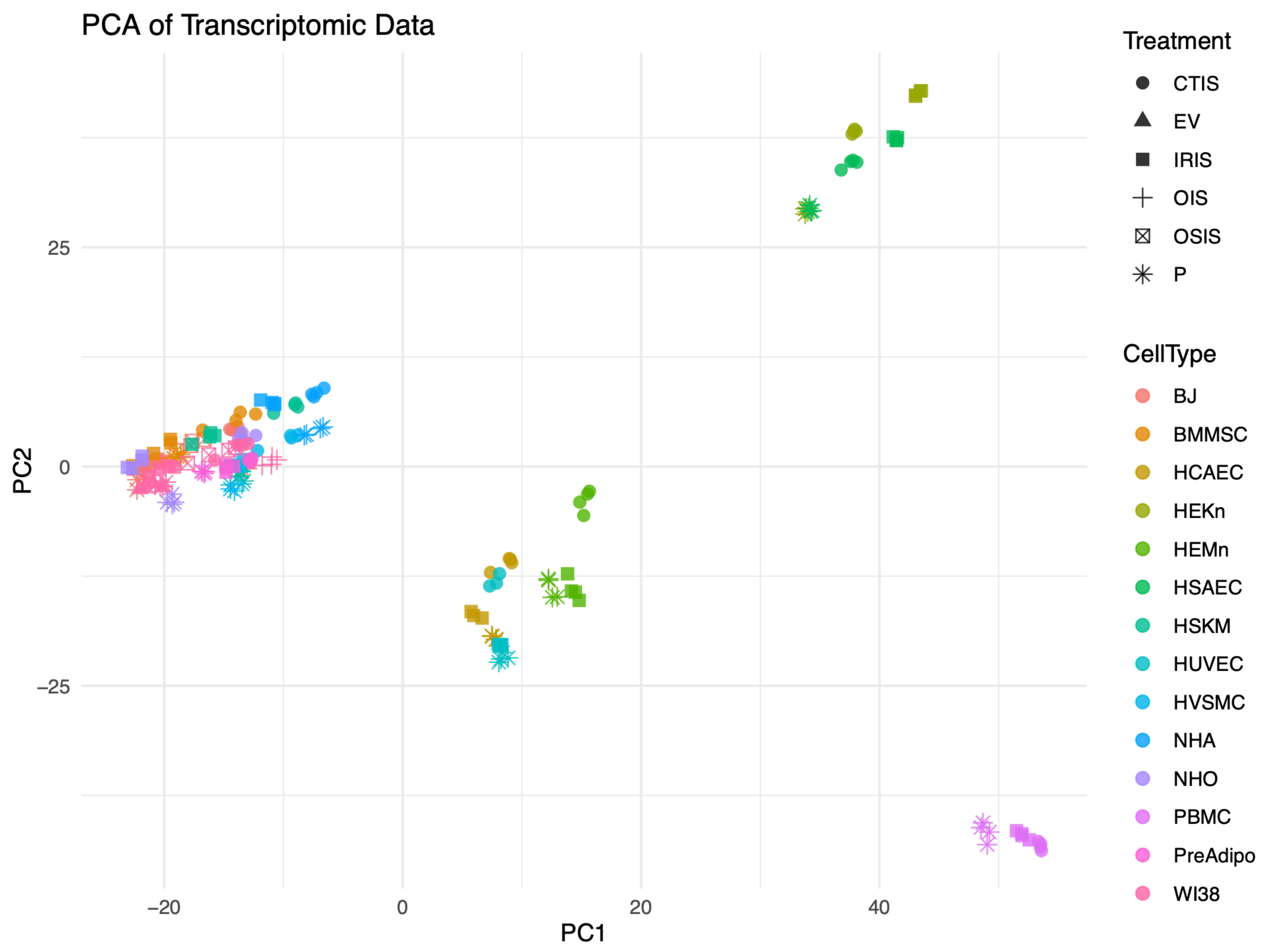
PCA of Transcriptomic Data
PC1
PC2
Treatment
CTIS
EV
IRIS
OIS
OSIS
P
CellType
BJ
BMMSC
HCAEC
HEKn
HEMn
HSAEC
HSKM
HUVEC
HVSMC
NHA
NHO
PBMC
PreAdipo
WI38

**A**

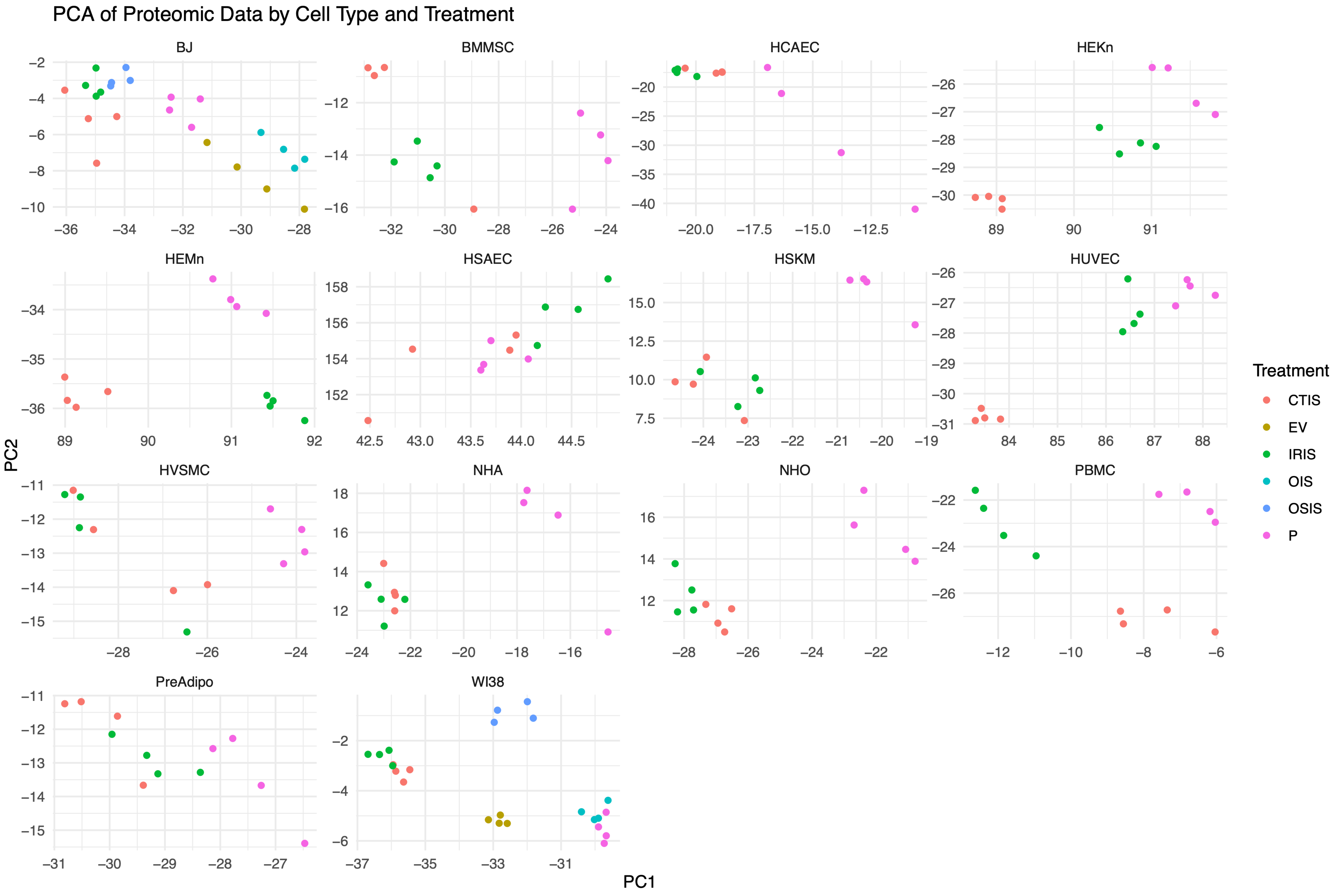
PCA of Proteomic Data by Cell Type and Treatment
BJ
BMMSC
HCAEC
HEKn
HEMn
HSAEC
HSKM
HUVEC
HVSMC
NHA
NHO
PBMC
PreAdipo
WI38
PC1
PC2
Treatment
CTIS
EV
IRIS
OIS
OSIS
P

**B**

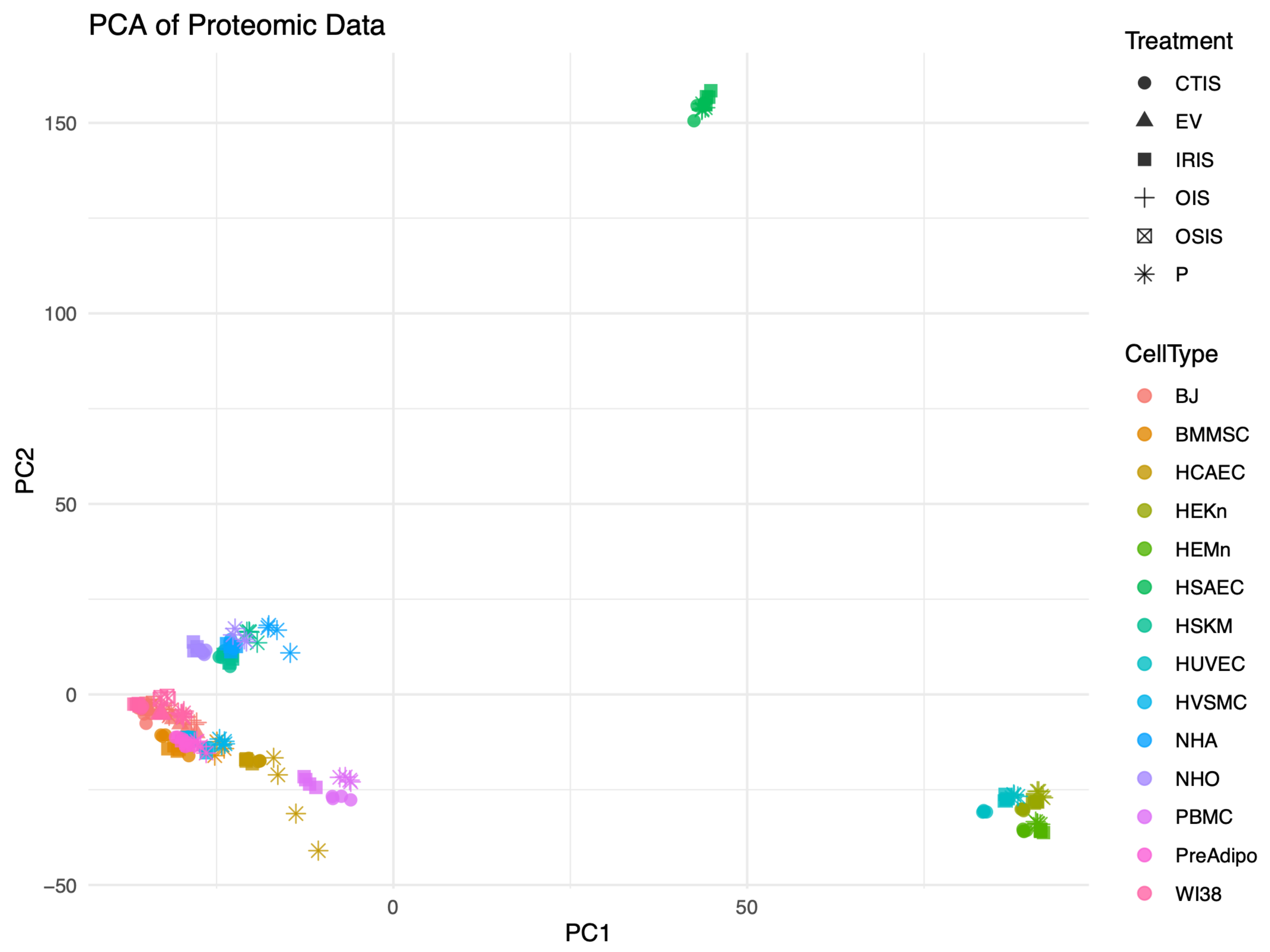
PCA of Proteomic Data
Treatment
CTIS
EV
IRIS
OIS
OSIS
P
CellType
BJ
BMMSC
HCAEC
HEKn
HEMn
HSAEC
HSKM
HUVEC
HVSMC
NHA
NHO
PBMC
PreAdipo
WI38
PC1
PC2

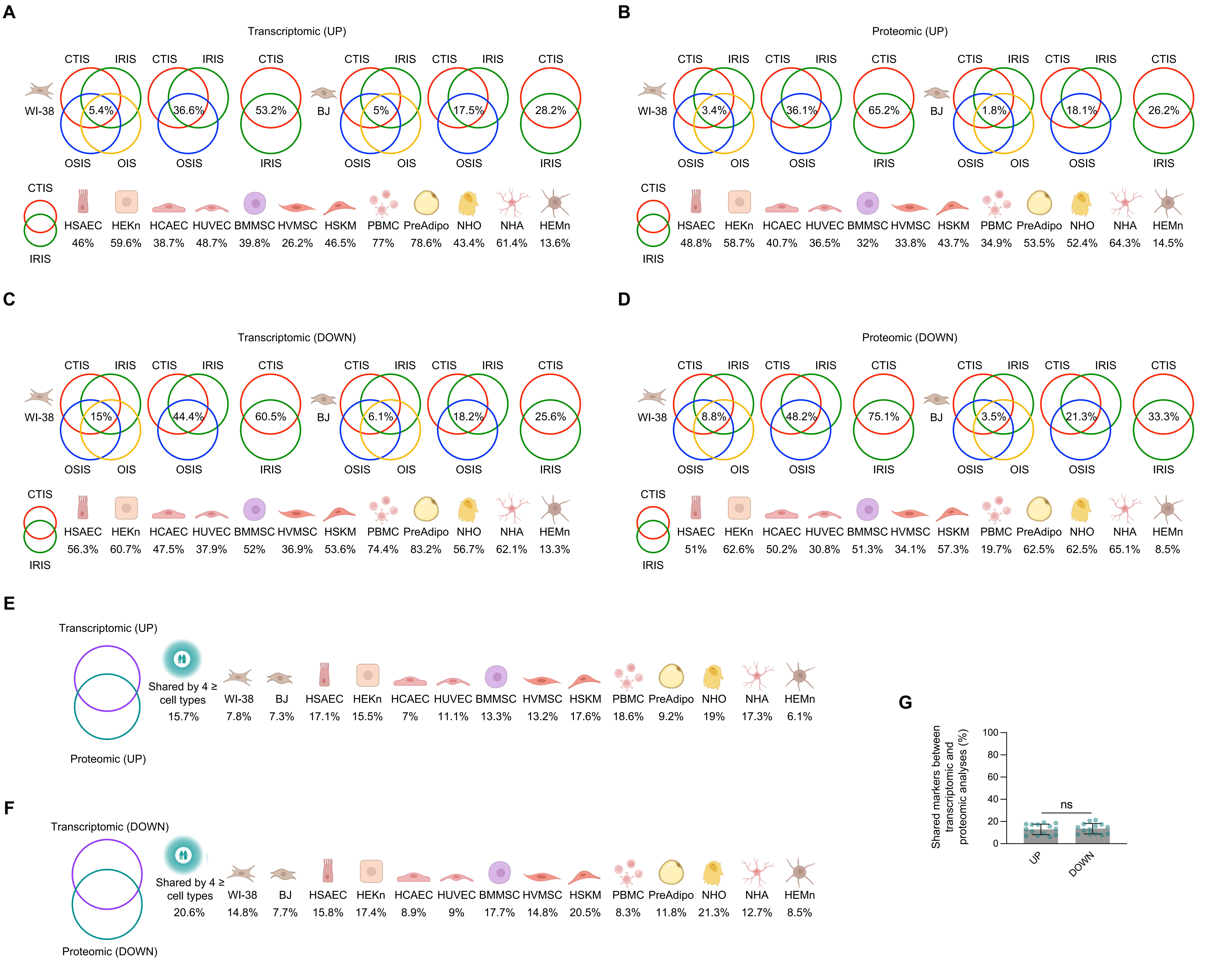
A
Transcriptomic (UP)
CTIS IRIS CTIS IRIS CTIS CTIS IRIS CTIS IRIS CTIS
WI-38 5.4% 36.6% 53.2% BJ 5% 17.5% 28.2%
OSIS OIS OSIS IRIS OSIS OIS OSIS IRIS
CTIS
IRIS
HSAEC 46% HEKn 59.6% HCAEC 38.7% HUVEC 48.7% BMMSC 39.8% HVMSC 26.2% HSKM 46.5% PBMC 77% PreAdipo 78.6% NHO 43.4% NHA 61.4% HEMn 13.6%
B
Proteomic (UP)
CTIS IRIS CTIS IRIS CTIS CTIS IRIS CTIS IRIS CTIS
WI-38 3.4% 36.1% 65.2% BJ 1.8% 18.1% 26.2%
OSIS OIS OSIS IRIS OSIS OIS OSIS IRIS
CTIS
IRIS
HSAEC 48.8% HEKn 58.7% HCAEC 40.7% HUVEC 36.5% BMMSC 32% HVMSC 33.8% HSKM 43.7% PBMC 34.9% PreAdipo 53.5% NHO 52.4% NHA 64.3% HEMn 14.5%
C
Transcriptomic (DOWN)
CTIS IRIS CTIS IRIS CTIS CTIS IRIS CTIS IRIS CTIS
WI-38 15% 44.4% 60.5% BJ 6.1% 18.2% 25.6%
OSIS OIS OSIS IRIS OSIS OIS OSIS IRIS
CTIS
IRIS
HSAEC 56.3% HEKn 60.7% HCAEC 47.5% HUVEC 37.9% BMMSC 52% HVMSC 36.9% HSKM 53.6% PBMC 74.4% PreAdipo 83.2% NHO 56.7% NHA 62.1% HEMn 13.3%
D
Proteomic (DOWN)
CTIS IRIS CTIS IRIS CTIS CTIS IRIS CTIS IRIS CTIS
WI-38 8.8% 48.2% 75.1% BJ 3.5% 21.3% 33.3%
OSIS OIS OSIS IRIS OSIS OIS OSIS IRIS
CTIS
IRIS
HSAEC 51% HEKn 62.6% HCAEC 50.2% HUVEC 30.8% BMMSC 51.3% HVMSC 34.1% HSKM 57.3% PBMC 19.7% PreAdipo 62.5% NHO 62.5% NHA 65.1% HEMn 8.5%
E
Transcriptomic (UP)
Proteomic (UP)
Shared by 4 ≥ cell types 15.7%
WI-38 7.8% BJ 7.3% HSAEC 17.1% HEKn 15.5% HCAEC 7% HUVEC 11.1% BMMSC 13.3% HVMSC 13.2% HSKM 17.6% PBMC 18.6% PreAdipo 9.2% NHO 19% NHA 17.3% HEMn 6.1%
F
Transcriptomic (DOWN)
Proteomic (DOWN)
Shared by 4 ≥ cell types 20.6%
WI-38 14.8% BJ 7.7% HSAEC 15.8% HEKn 17.4% HCAEC 8.9% HUVEC 9% BMMSC 17.7% HVMSC 14.8% HSKM 20.5% PBMC 8.3% PreAdipo 11.8% NHO 21.3% NHA 12.7% HEMn 8.5%
G
Shared markers between transcriptomic and proteomic analyses (%)
100
80
60
40
20
0
ns
UP
DOWN

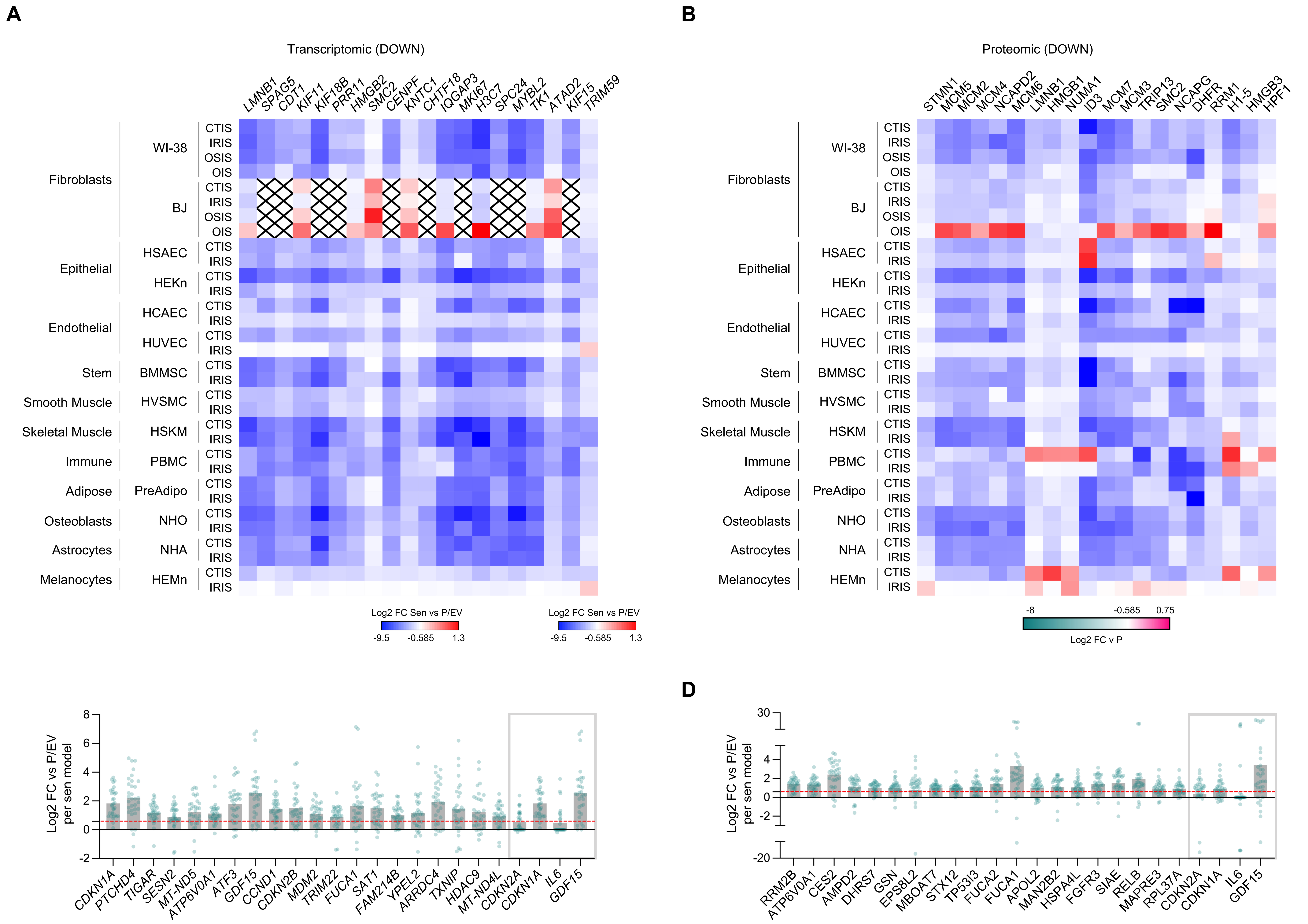
A
Transcriptomic (DOWN)
B
Proteomic (DOWN)
Fibroblasts
WI-38
BJ
Epithelial
HSAEC
HEKn
Endothelial
HCAEC
HUVEC
Stem
BMMSC
Smooth Muscle
HVSMC
Skeletal Muscle
HSKM
Immune
PBMC
Adipose
PreAdipo
Osteoblasts
NHO
Astrocytes
NHA
Melanocytes
HEMn
CTIS
IRIS
OSIS
OIS
Log2 FC Sen vs P/EV
-9.5
-0.585
1.3
Log2 FC v P
-8
0.75
C
D
Log2 FC vs P/EV per sen model

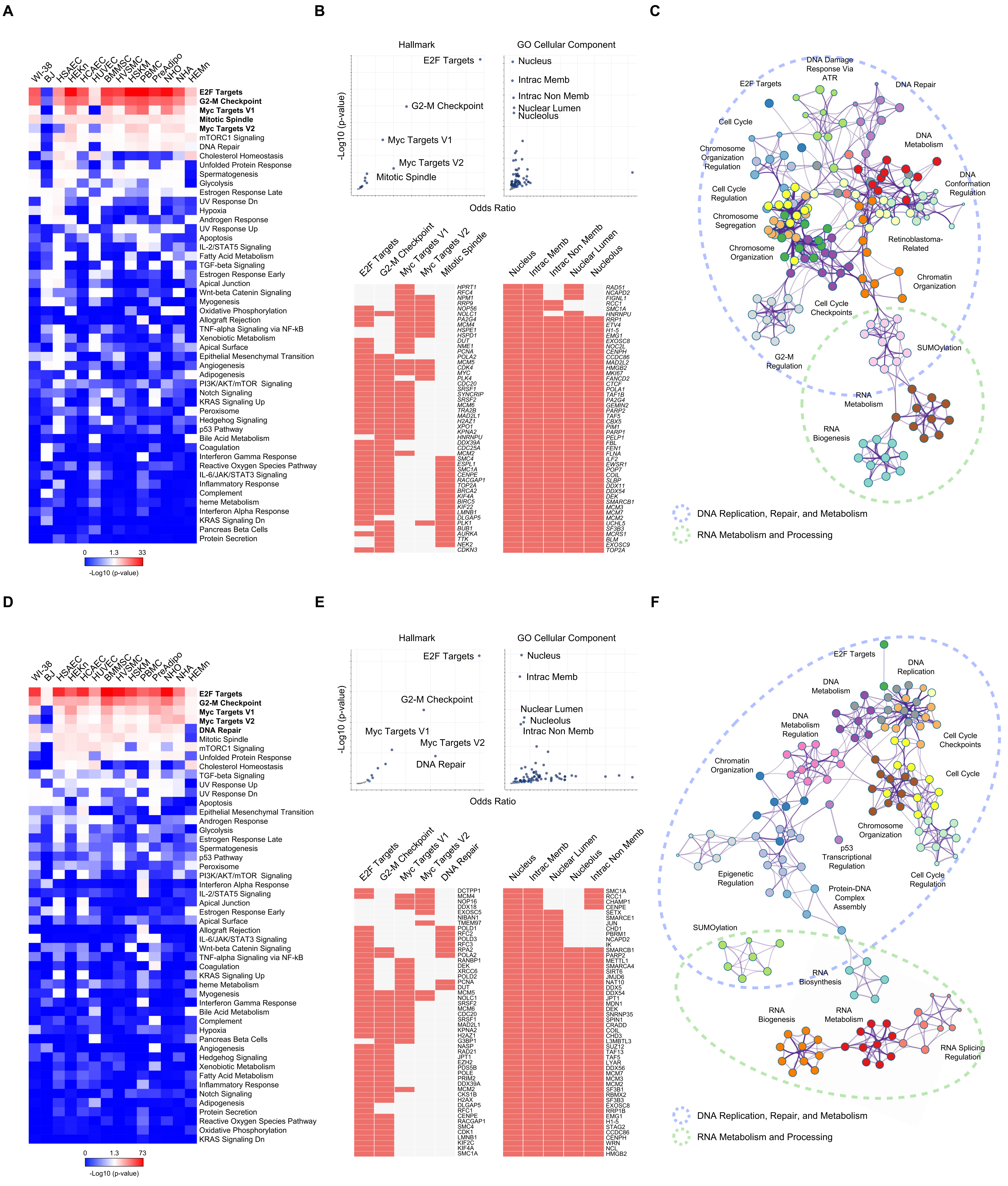
A
WI-38
BJ
HSAEC
HEKn
HCAEC
HUVEC
BMMSC
HVSMC
HSKM
PBMC
PreAdipo
NHO
NHA
HEMn
E2F Targets
G2-M Checkpoint
Myc Targets V1
Mitotic Spindle
Myc Targets V2
mTORC1 Signaling
DNA Repair
Cholesterol Homeostasis
Unfolded Protein Response
Spermatogenesis
Glycolysis
Estrogen Response Late
UV Response Dn
Hypoxia
Androgen Response
UV Response Up
Apoptosis
IL-2/STAT5 Signaling
Fatty Acid Metabolism
TGF-beta Signaling
Estrogen Response Early
Apical Junction
Wnt-beta Catenin Signaling
Myogenesis
Oxidative Phosphorylation
Allograft Rejection
TNF-alpha Signaling via NF-kB
Xenobiotic Metabolism
Apical Surface
Epithelial Mesenchymal Transition
Angiogenesis
Adipogenesis
PI3K/AKT/mTOR Signaling
Notch Signaling
KRAS Signaling Up
Peroxisome
Hedgehog Signaling
p53 Pathway
Bile Acid Metabolism
Coagulation
Interferon Gamma Response
Reactive Oxygen Species Pathway
IL-6/JAK/STAT3 Signaling
Inflammatory Response
Complement
heme Metabolism
Interferon Alpha Response
KRAS Signaling Dn
Pancreas Beta Cells
Protein Secretion
0 1.3 33
-Log10 (p-value)
B
Hallmark
GO Cellular Component
E2F Targets
G2-M Checkpoint
Myc Targets V1
Myc Targets V2
Mitotic Spindle
Nucleus
Intrac Memb
Intrac Non Memb
Nuclear Lumen
Nucleolus
-Log10 (p-value)
Odds Ratio
C
DNA Damage Response Via ATR
E2F Targets
DNA Repair
Cell Cycle
DNA Metabolism
Chromosome Organization Regulation
DNA Conformation Regulation
Cell Cycle Regulation
Chromosome Segregation
Retinoblastoma-Related
Chromosome Organization
Chromatin Organization
Cell Cycle Checkpoints
SUMOylation
G2-M Regulation
RNA Metabolism
RNA Biogenesis
DNA Replication, Repair, and Metabolism
RNA Metabolism and Processing
D
WI-38
BJ
HSAEC
HEKn
HCAEC
HUVEC
BMMSC
HVSMC
HSKM
PBMC
PreAdipo
NHO
NHA
HEMn
E2F Targets
G2-M Checkpoint
Myc Targets V1
Myc Targets V2
DNA Repair
Mitotic Spindle
mTORC1 Signaling
Unfolded Protein Response
Cholesterol Homeostasis
TGF-beta Signaling
UV Response Up
UV Response Dn
Apoptosis
Epithelial Mesenchymal Transition
Androgen Response
Glycolysis
Estrogen Response Late
Spermatogenesis
p53 Pathway
Peroxisome
PI3K/AKT/mTOR Signaling
Interferon Alpha Response
IL-2/STAT5 Signaling
Apical Junction
Estrogen Response Early
Apical Surface
Allograft Rejection
IL-6/JAK/STAT3 Signaling
Wnt-beta Catenin Signaling
TNF-alpha Signaling via NF-kB
Coagulation
KRAS Signaling Up
heme Metabolism
Myogenesis
Interferon Gamma Response
Bile Acid Metabolism
Complement
Hypoxia
Pancreas Beta Cells
Angiogenesis
Hedgehog Signaling
Xenobiotic Metabolism
Fatty Acid Metabolism
Inflammatory Response
Notch Signaling
Adipogenesis
Protein Secretion
Reactive Oxygen Species Pathway
Oxidative Phosphorylation
KRAS Signaling Dn
0 1.3 73
-Log10 (p-value)
E
Hallmark
GO Cellular Component
E2F Targets
G2-M Checkpoint
Myc Targets V1
Myc Targets V2
DNA Repair
Nucleus
Intrac Memb
Nuclear Lumen
Nucleolus
Intrac Non Memb
-Log10 (p-value)
Odds Ratio
F
E2F Targets
DNA Replication
DNA Metabolism
DNA Metabolism Regulation
Cell Cycle Checkpoints
Chromatin Organization
Cell Cycle
Chromosome Organization
p53 Transcriptional Regulation
Epigenetic Regulation
Cell Cycle Regulation
Protein-DNA Complex Assembly
SUMOylation
RNA Biosynthesis
RNA Biogenesis
RNA Metabolism
RNA Splicing Regulation
DNA Replication, Repair, and Metabolism
RNA Metabolism and Processing

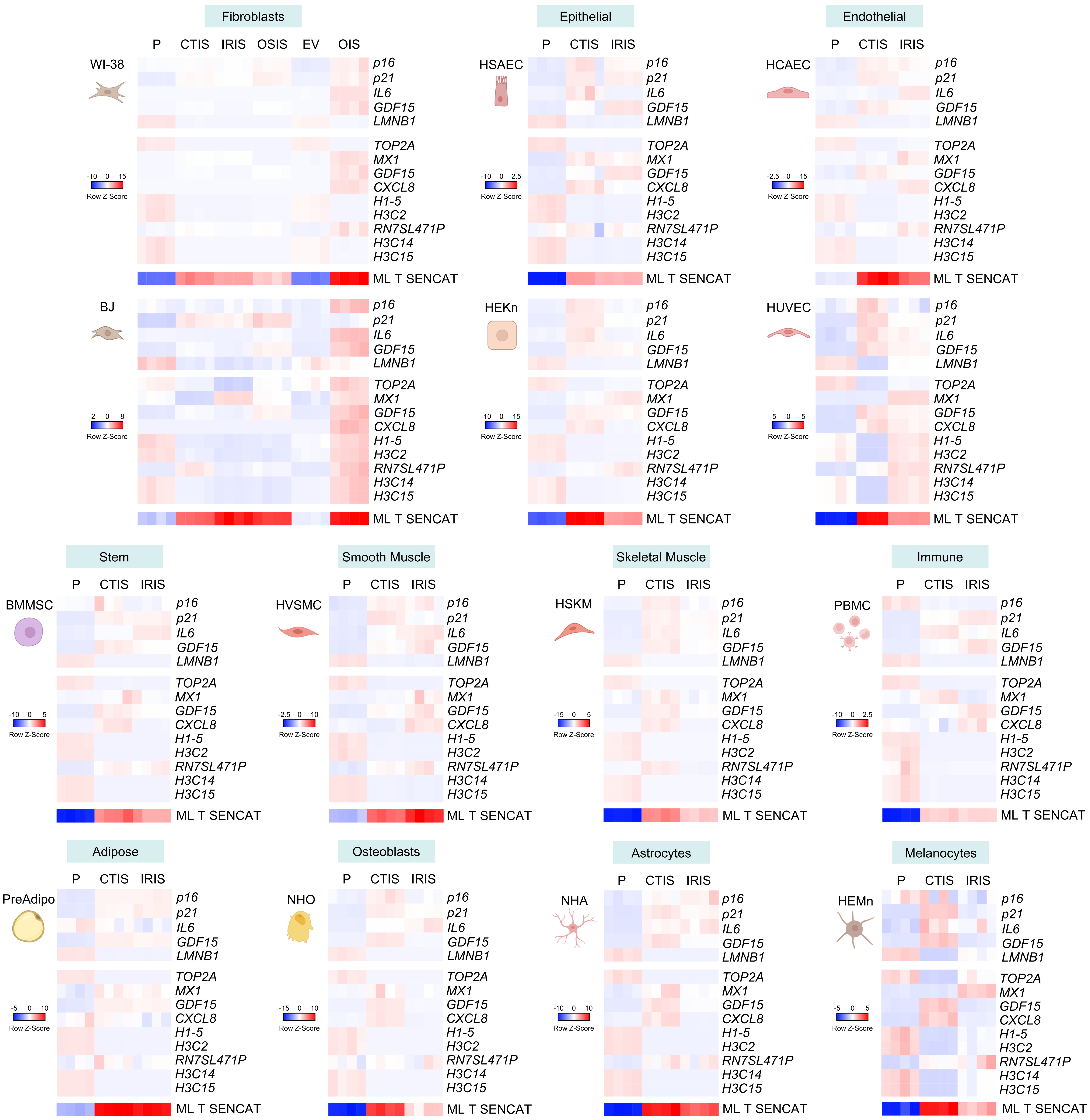
Fibroblasts
P CTIS IRIS OSIS EV OIS
WI-38
BJ
Epithelial
P CTIS IRIS
HSAEC
HEKn
Endothelial
P CTIS IRIS
HCAEC
HUVEC
Stem
BMMSC
Smooth Muscle
HVSMC
Skeletal Muscle
HSKM
Immune
PBMC
Adipose
PreAdipo
Osteoblasts
NHO
Astrocytes
NHA
Melanocytes
HEMn
p16
p21
IL6
GDF15
LMNB1
TOP2A
MX1
GDF15
CXCL8
H1-5
H3C2
RN7SL471P
H3C14
H3C15
ML T SENCAT
Row Z-Score

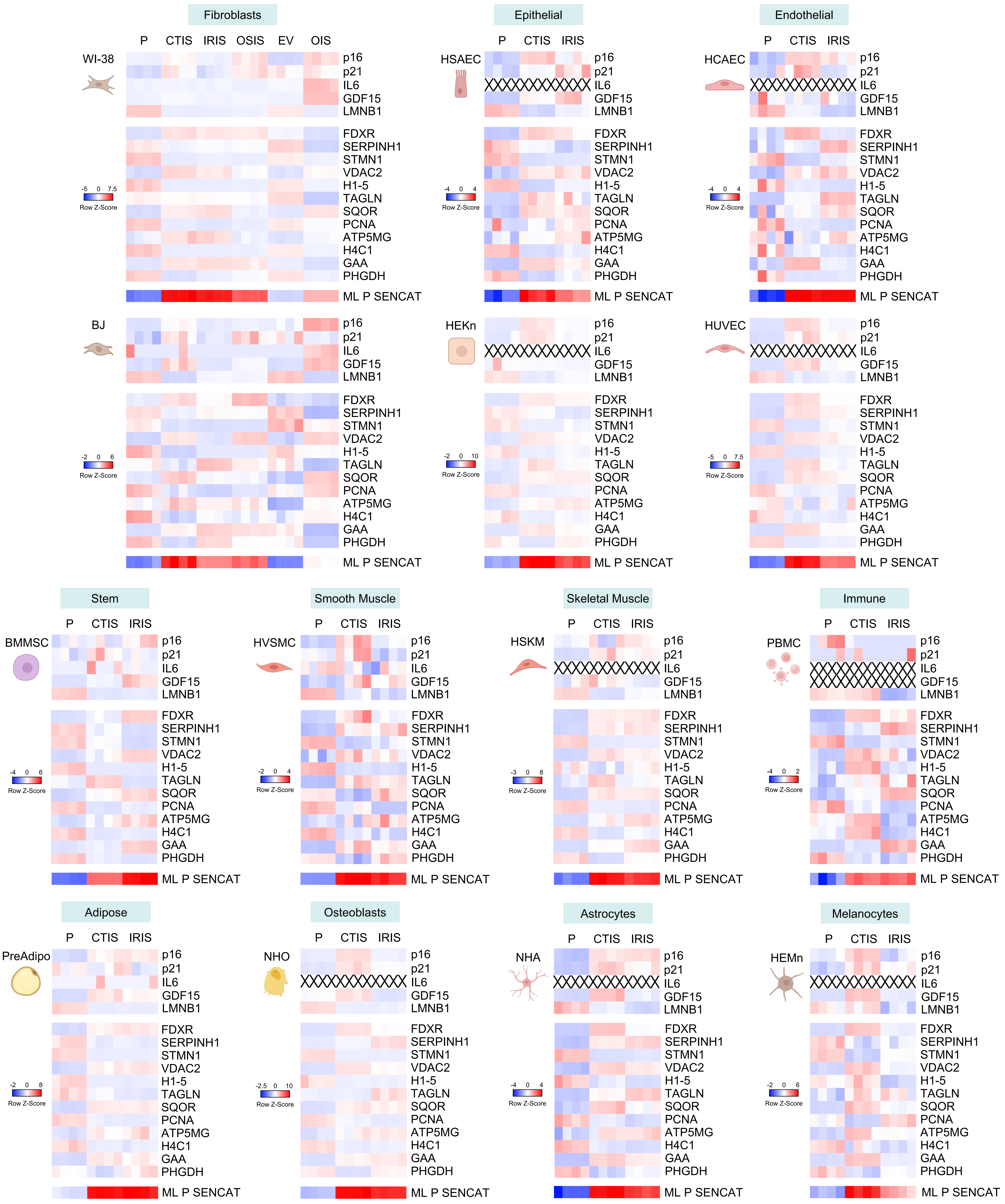
Fibroblasts
Epithelial
Endothelial
P CTIS IRIS OSIS EV OIS
P CTIS IRIS
WI-38
HSAEC
HCAEC
BJ
HEKn
HUVEC
p16
p21
IL6
GDF15
LMNB1
FDXR
SERPINH1
STMN1
VDAC2
H1-5
TAGLN
SQOR
PCNA
ATP5MG
H4C1
GAA
PHGDH
ML P SENCAT
-5 0 7.5
-4 0 4
-2 0 6
-2 0 10
Row Z-Score
Stem
Smooth Muscle
Skeletal Muscle
Immune
BMMSC
HVSMC
HSKM
PBMC
-4 0 6
-2 0 4
-3 0 8
-4 0 2
Adipose
Osteoblasts
Astrocytes
Melanocytes
PreAdipo
NHO
NHA
HEMn
-2 0 8
-2.5 0 10

A

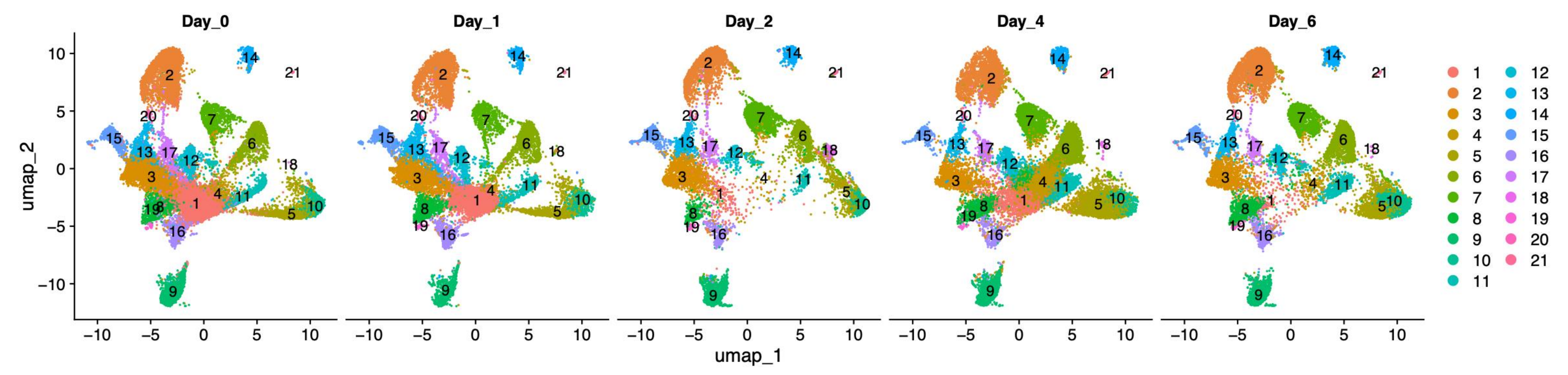


B

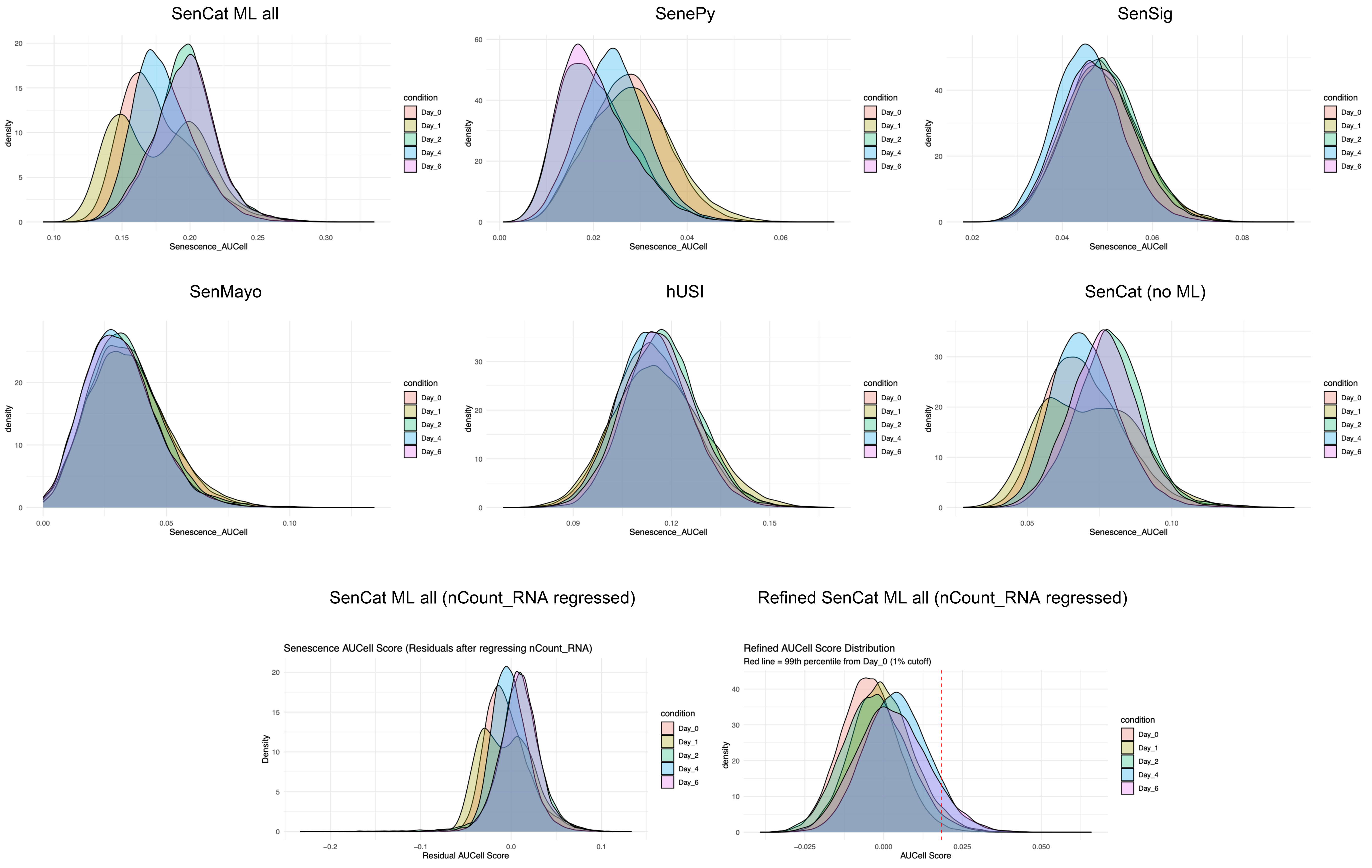


C

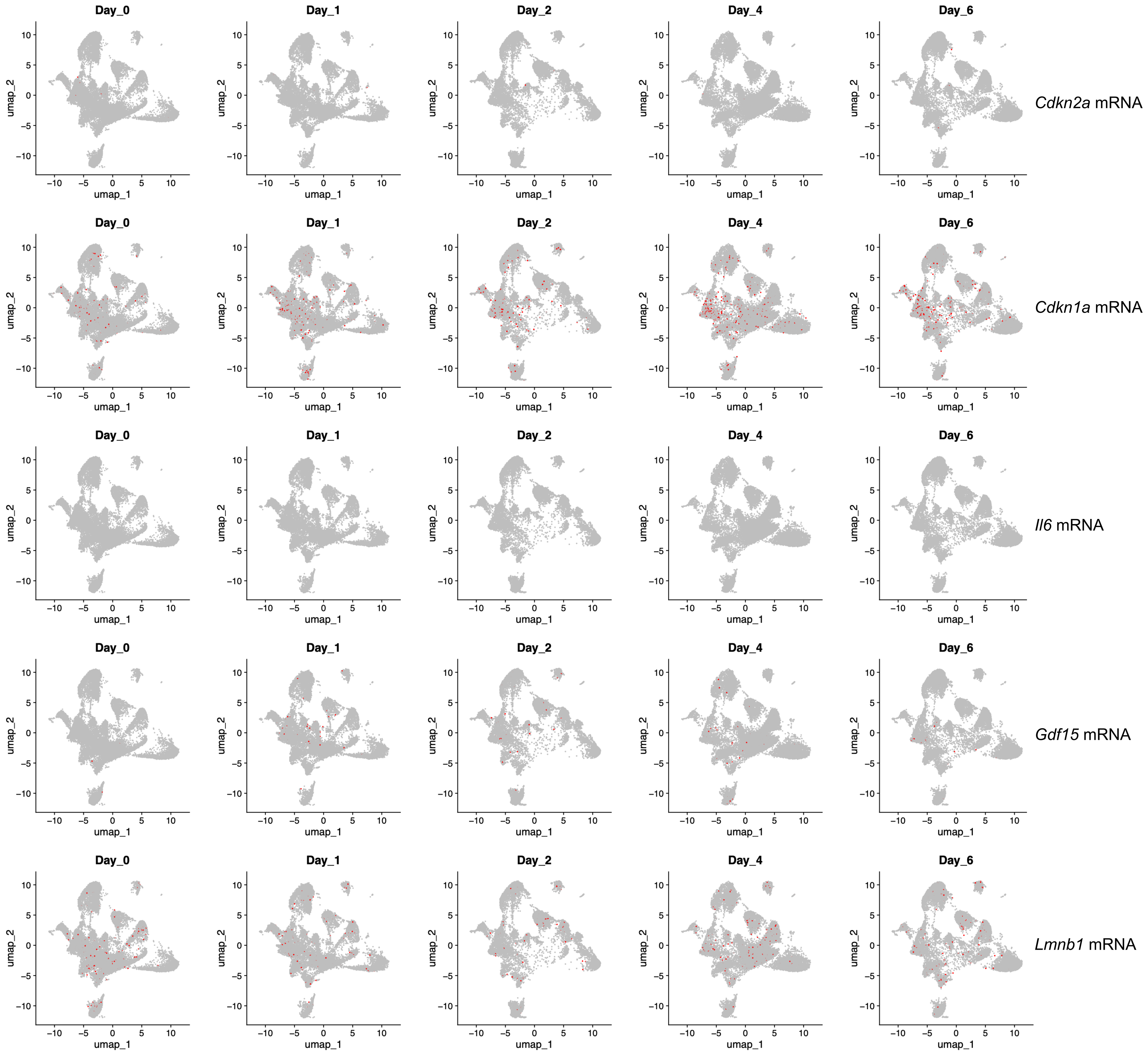


D

| Condition | Total cells | Senescent cells | Percentage senescent |
|---|---|---|---|
| Day_0 | 30783 | 308 | 1 |
| Day_1 | 19916 | 775 | 3.89 |
| Day_2 | 13448 | 374 | 2.78 |
| Day_4 | 29505 | 2646 | 8.97 |
| Day_6 | 16716 | 1517 | 9.08 |

A

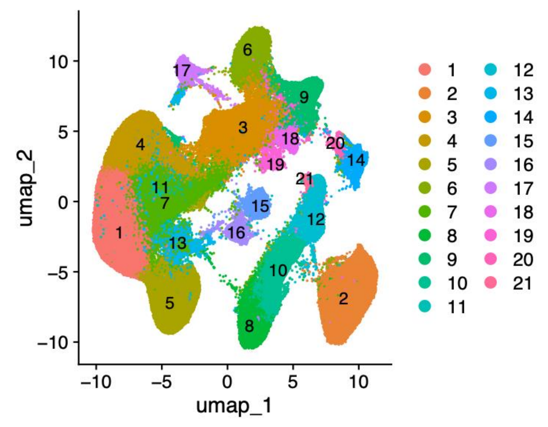

B

| Cluster | Cell Type |
|---|---|
| 1 | Epithelial (Proximal Tubule) |
| 2 | Epithelial (Distal Convoluted Tube) |
| 3 | Epithelial (Proximal Tubule) |
| 4 | Epithelial (Proximal Tubule) |
| 5 | Epithelial (Proximal Tubule) |
| 6 | Epithelial (Proximal Tubule) |
| 7 | Epithelial (Proximal Tubule) |
| 8 | Epithelial (Distal Convoluted Tube) |
| 9 | Podocyte |
| 10 | Epithelial (Distal Convoluted Tube) |
| 11 | Epithelial (Proximal Tubule) |
| 12 | Epithelial (Collecting Duct) |
| 13 | Epithelial (Proximal Tubule) |
| 14 | Epithelial (Collecting Duct) |
| 15 | Epithelial (Collecting Duct) |
| 16 | Epithelial (Collecting Duct) |
| 17 | Podocyte |
| 18 | Smooth Muscle Cell |
| 19 | Macrophages |
| 20 | Fibroblast |
| 21 | Epithelial (Distal Convoluted Tube) |

C

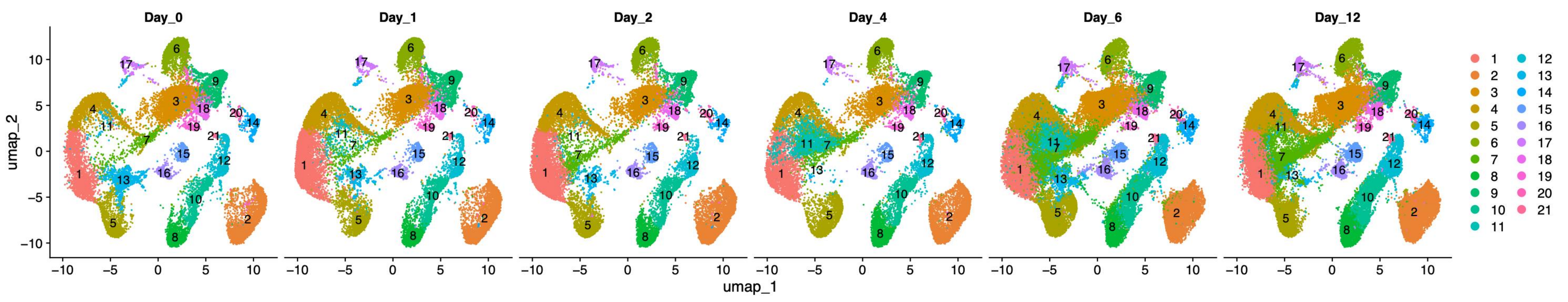


D

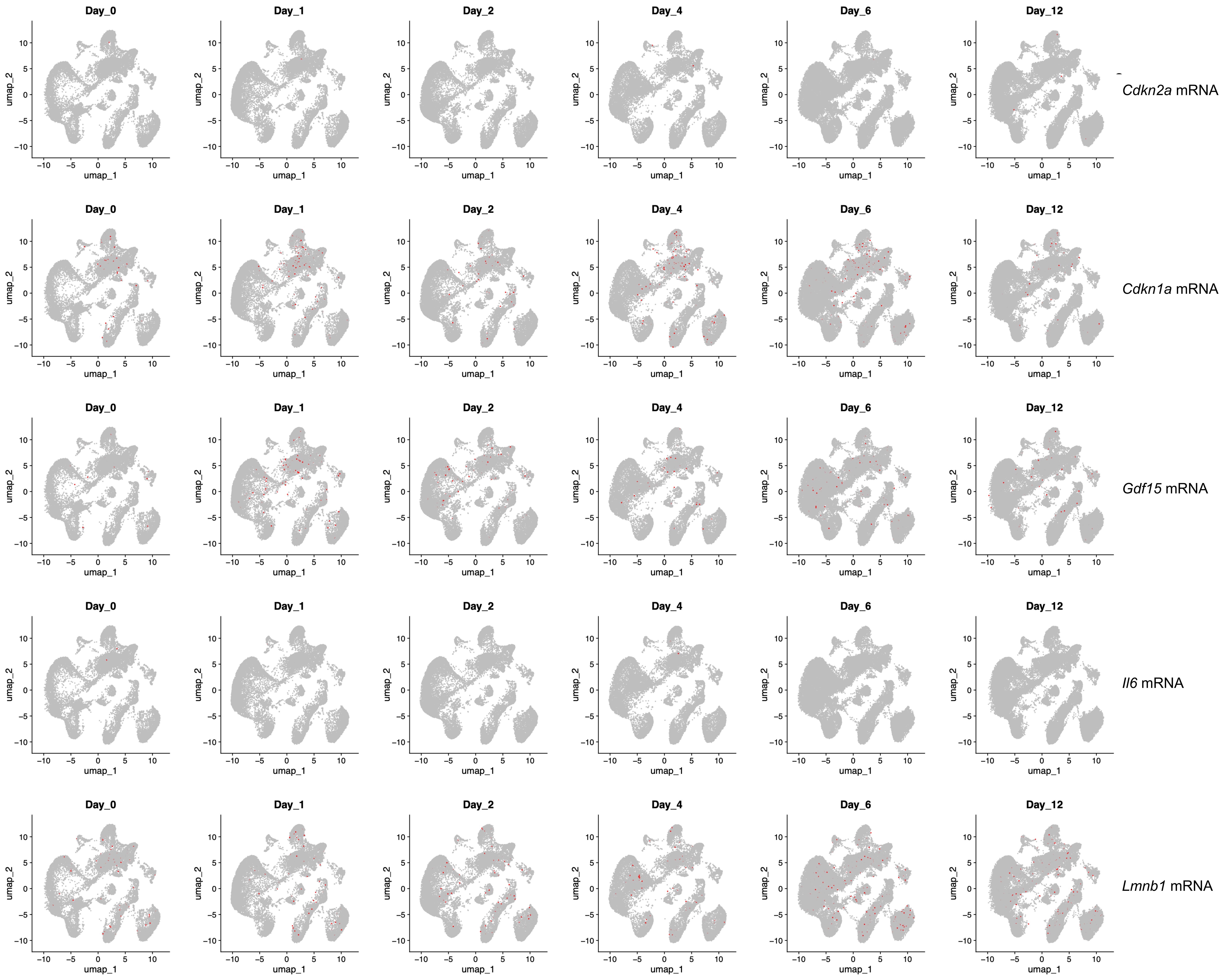


E

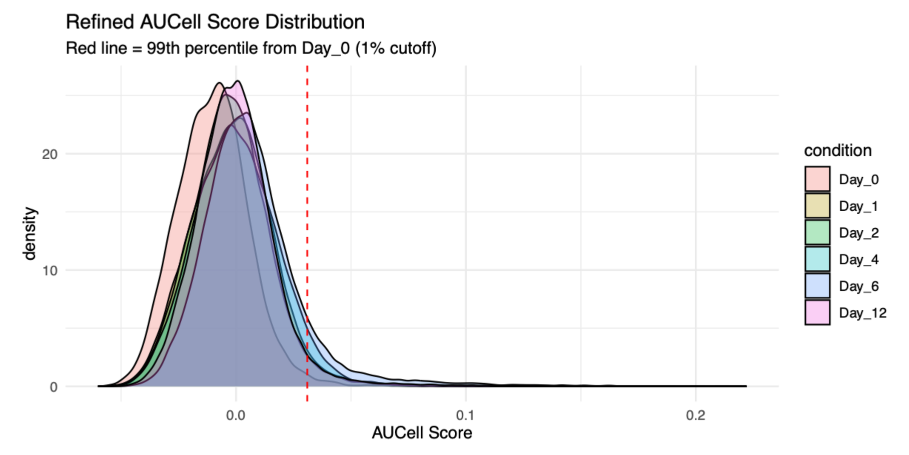


F

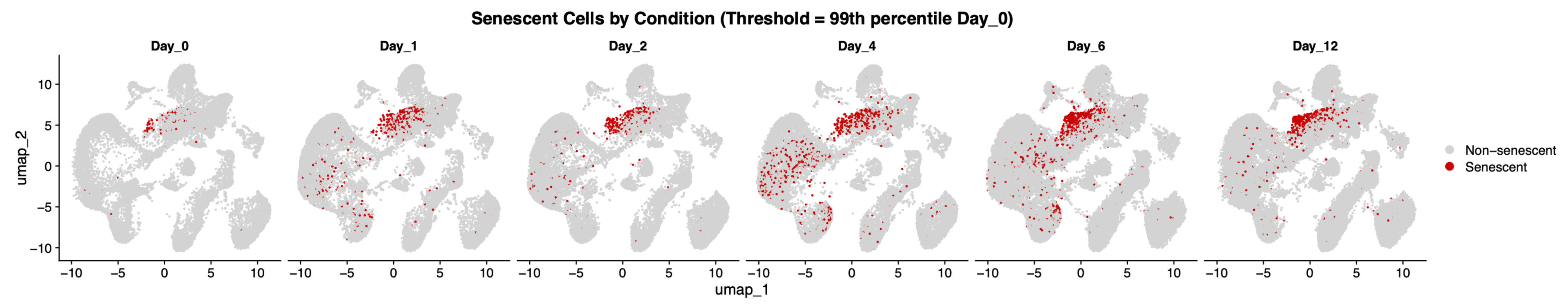


G

| condition | cells | senescent_cells | percent_senescent |
|---|---|---|---|
| Day_0 | 22049 | 221 | 1 |
| Day_1 | 25351 | 863 | 3.4 |
| Day_2 | 22582 | 716 | 3.17 |
| Day_4 | 21961 | 1083 | 4.93 |
| Day_6 | 43794 | 4292 | 9.8 |
| Day_12 | 49446 | 1881 | 3.8 |

H

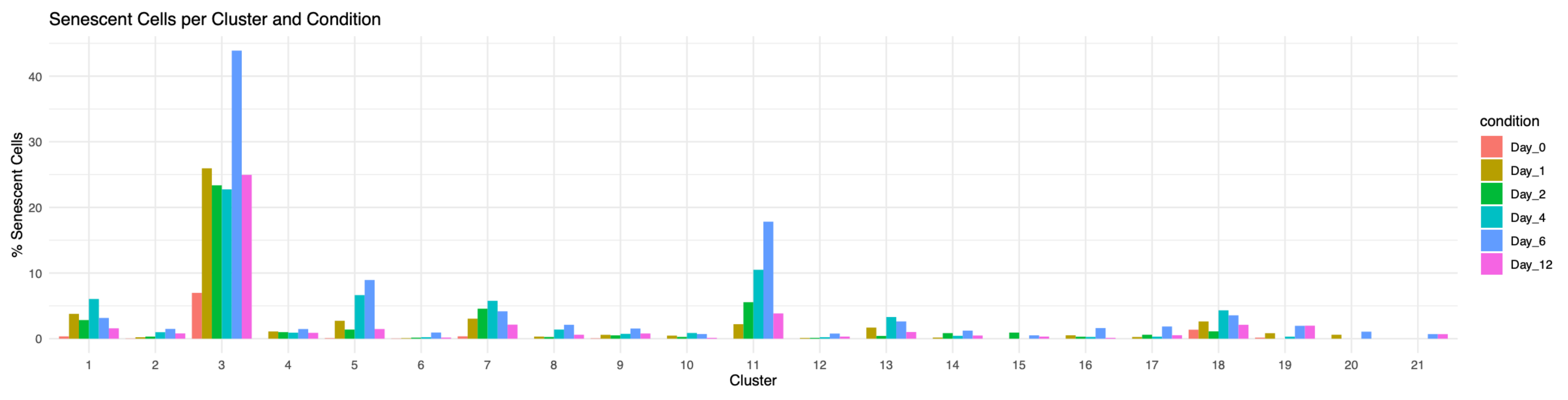

A

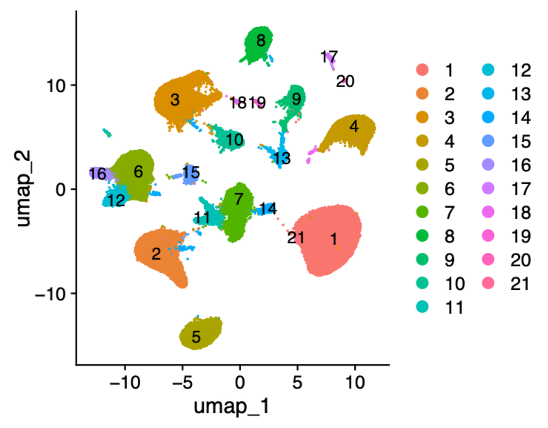


B

| Cluster | Cell type |
|---|---|
| 1 | Epithelial (AT2) |
| 2 | B Cell |
| 3 | Endothelial |
| 4 | Epithelial (Clara cell) |
| 5 | Macrophage |
| 6 | T Cell |
| 7 | Monocyte |
| 8 | Epithelial (AT1) |
| 9 | Smooth Muscle Cell |
| 10 | Endothelial |
| 11 | Monocyte |
| 12 | T Cell |
| 13 | Fibroblast |
| 14 | Neutrophil |
| 15 | B Cell |
| 16 | T Cell |
| 17 | Epithelial (Ciliated cell) |
| 18 | Fibroblast |
| 19 | Epithelial (AT1) |
| 20 | Fibroblast |
| 21 | Epithelial (AT2) |

C

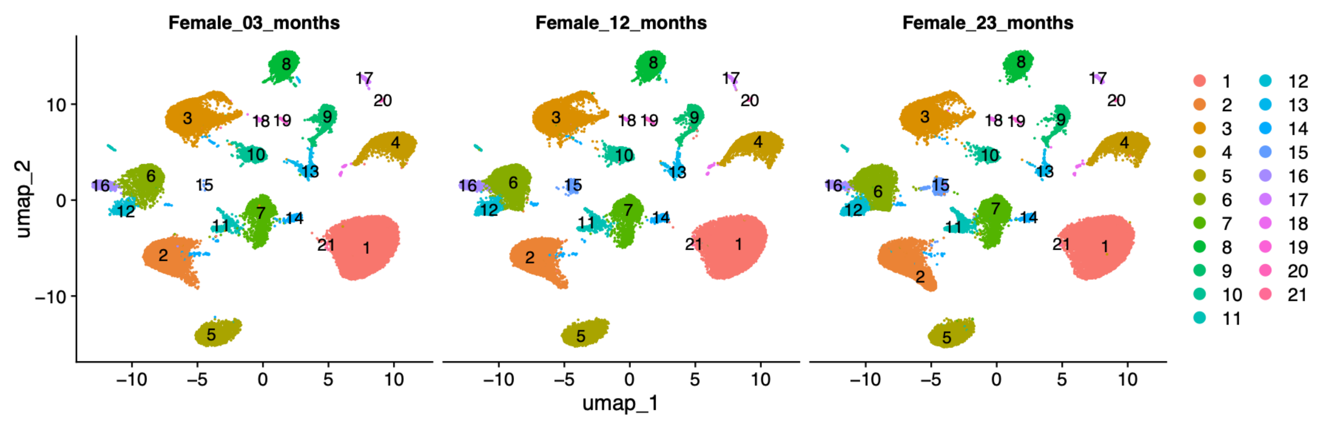


D

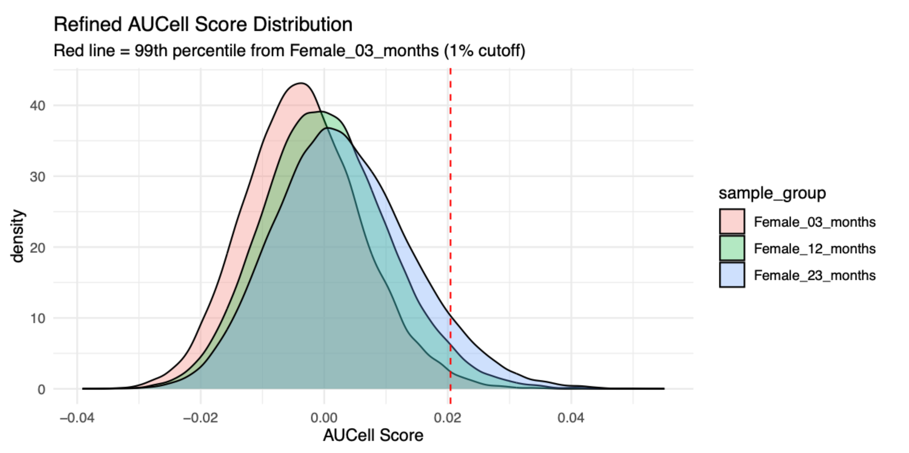


E

| sample_gro | total_cells | senescent_cells | percent_senescent |
|---|---|---|---|
| Female_03_ | 40375 | 404 | 1 |
| Female_12_ | 40212 | 1301 | 3.24 |
| Female_23_ | 36849 | 2572 | 6.98 |

F

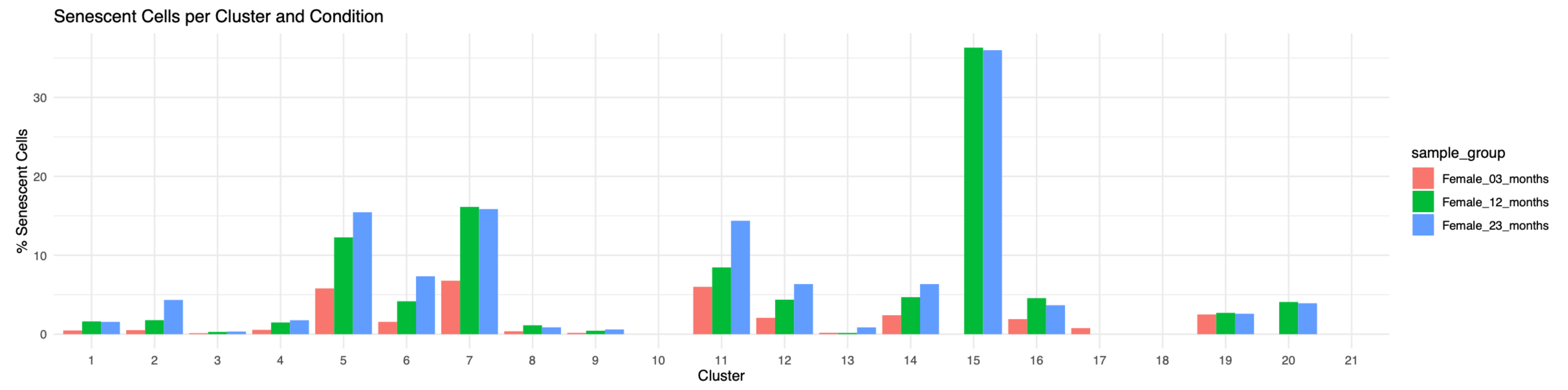


G

| interacting_pair | gene_a | gene_b | Senescent\|Neighbor |
|---|---|---|---|
| FGF10_FGFR2 | FGF10 | FGFR2 | 1,502 |
| EFNA5_EPHA7 | EFNA5 | EPHA7 | 713 |
| SEMA3A_NRP1 | SEMA3A | NRP1 | 604 |
| SEMA3A_PlexinA4_complex1 | SEMA3A | | 503 |
| IGF1_IGF1R | IGF1 | IGF1R | 471 |
| SEMA6A_PlexinA2_complex1 | SEMA6A | | 410 |
| THBS1_CD36 | THBS1 | CD36 | 396 |
| TNC_integrin_a9b1_complex | TNC | | 378 |
| NRXN1_DAG1 | NRXN1 | DAG1 | 285 |
| TGFB2_TGFbeta_receptor2 | TGFB2 | | 202 |
