## Supplementary material for "SenCat: Cataloging human cell senescence through multiomic profiling of multiple senescent primary cell types": Legends to Supplementary Tables

**LEGENDS FOR SUPPLEMENTARY TABLES**

**Supplementary Table S1. Transcriptomic Results.**
Raw transcriptomic expression matrix for all samples across cell types and conditions. Rows correspond to Ensembl gene IDs and gene symbols. Columns represent RNA-seq counts (normalized counts) for each sample.

**Supplementary Table S2. Proteomic Results.**
Protein abundance matrix for all samples and conditions derived from mass spectrometry analysis. Rows include gene symbols and UniProt IDs. Columns represent normalized protein intensities for each sample, grouped by cell type and condition.

**Supplementary Table S3. Markers increased by transcriptomic analysis.**
List of transcriptomic senescence markers upregulated in different conditions and cell types. Columns represent cell types and specific senescence comparisons (e.g., CTIS compared to P, IRIS compared to P). Each cell contains transcripts significantly upregulated (log2FC > 0.585, p-val < 0.05) under the specified condition.

**Supplementary Table S4. Markers reduced by transcriptomic analysis.**
List of transcriptomic senescence markers downregulated in different conditions and cell types. Columns represent cell types and specific senescence comparisons (e.g., CTIS compared to P, IRIS compared to P). Each cell contains transcripts significantly downregulated (log2FC < -0.585, p-val < 0.05) under the specified condition.

**Supplementary Table S5. Markers increased by proteomic analysis.**

List of proteomic senescence markers upregulated in different conditions and cell types. Columns represent cell types and specific senescence comparisons (e.g., CTIS compared to P, IRIS compared to P). Each cell contains proteins significantly upregulated (log2FC > 0.585, p-val < 0.05) under the specified condition.

**Supplementary Table S6. Markers reduced by proteomic analysis.**
List of proteomic senescence markers downregulated in different conditions and cell types. Columns represent cell types and specific senescence comparisons (e.g., CTIS compared to P, IRIS compared to P). Each cell contains proteins significantly downregulated (log2FC < -0.585, p-val < 0.05) under the specified condition.

**Supplementary Table S7. Machine learning–derived transcriptomic markers and coefficients.**

List of transcriptomic senescence marker genes derived using machine learning (ML) models trained across all cell types. Each row includes the Ensembl gene ID, gene symbol, and the model coefficient indicating direction and importance in senescence classification.

**Supplementary Table S8. Senescence markers from other publications.**
Reference list of senescence markers compiled from previous studies, including SenMayo, SenePy, SenSig (filtered for FC > 1.5), and hUSI datasets. Used for benchmarking and comparison with the SenCat-derived markers.

**Supplementary Table S9. Conditions of senescence induction.**
Experimental conditions used to induce senescence in the specified primary human cell types. Columns include the cell type, senescence trigger (e.g., doxorubicin, ionizing radiation, etoposide), treatment details (dose and duration), and number of days after treatment at which cells were profiled.
